# MVPAlab: A Machine Learning decoding toolbox for multidimensional electroencephalography data

**DOI:** 10.1101/2021.06.24.449693

**Authors:** David López-García, Jose M.G. Peñalver, Juan M. Górriz, María Ruz

**Affiliations:** Mind, Brain and Behavior Research Center. University of Granada, Spain; Data Science & Computational Intelligence Institute, University of Granada, Spain; Mind, Brain and Behavior Research Center. Department of Experimental Psychology. University of Granada, Spain

**Keywords:** machine learning, classification, cross-classification, decoding, cross-validation, multivariate pattern analysis, mvpa, EEG, MEG, MVPAlab toolbox

## Abstract

**Background and Objective:** The study of brain function has recently expanded from classical univariate to multivariate analyses. These multivariate, machine learning-based algorithms afford neuroscientists extracting more detailed and richer information from the data. However, the implementation of these procedures is usually challenging, especially for researchers with no coding experience. To address this problem, we have developed MVPAlab, a MATLAB-based, flexible decoding toolbox for multidimensional electroencephalography and magnetoencephalography data.

**Methods:** The MVPAlab Toolbox implements several machine learning algorithms to compute multivariate pattern analyses, cross-classification, temporal generalization matrices and feature and frequency contribution analyses. It also provides access to an extensive set of preprocessing routines for, among others, data normalization, data smoothing, dimensionality reduction and supertrial generation. To draw statistical inferences at the group level, MVPAlab includes a non-parametric cluster-based permutation approach.

**Results:** A sample electroencephalography dataset was compiled to test all the MVPAlab main functionalities. Significant clusters *(p<0.01)* were found for the proposed decoding analyses and different configurations, proving the software capability for discriminating between different experimental conditions.

**Conclusions:** This toolbox has been designed to include an easy-to-use and intuitive graphic user interface and data representation software, which makes MVPAlab a very convenient tool for users with few or no previous coding experience. In addition, MVPAlab is not for beginners only, as it implements several high and low-level routines allowing more experienced users to design their own projects in a highly flexible manner.

## 1. INTRODUCTION

Historically, the study of brain function employing electroencephalography (EEG) data has relied on classical univariate analyses of amplitudes and delays of different peaks of the average of several evoked EEG recordings, commonly called Event-Related Potentials (ERPs). The constant development of science and technology in past decades has allowed researchers and engineers to develop and apply more advanced signal processing techniques, such as time/frequency analyses, phase clustering, Independent Component Analysis (ICA) decompositions[1,2], and others. These techniques have been implemented in excellent analysis and preprocessing tools, such as EEGLAB [3], ERPLAB [4] or Fieldtrip [5], enabling researchers to develop a myriad of studies in a wide range of areas.

More recently, newer Machine Learning-based algorithms (ML), in conjunction with advanced neuroimaging techniques, such as functional Magnetic Resonance Imaging (fMRI) or Magnetoencephalography (MEG), have gained popularity in neuroscience. This trend started with studies by Haxby and Norman [6–8], and other reference contributions [9–14], which opened novel avenues of research on brain function. For years, ML models have been also successfully employed in medical imaging, mainly in the area of computer-aided diagnosis[15]. To mention just a few examples, the use of different ML approaches is mainstream in the study and detection of several neurological diseases, such as Parkinson[16–18], Alzheimer[19–21], Autism[22–24], or sleep disorders[25–27]. Even the recently spread COVID-19 can be successfully diagnosed using Artificial Intelligence (AI) in chest radiographies, according to preliminary studies[28–30]. However, the recent growth of ML models is not limited to neuroscience or medical applications but is present in a huge range of scientific disciplines in a cross-cutting basis.

### 1.1 Related work

Multivariate Pattern Analysis (MVPA) usually encompasses a set of supervised learning algorithms, which provide a theoretically elegant, computationally efficient, and very effective solution in many practical pattern recognition scenarios. One of the most remarkable advantages of these multivariate approaches over univariate ones is its sensitivity in unveiling subtle changes in activations associated with specific information content in brain patterns. Several MVPA toolboxes, such as SPM [31], The Decoding Toolbox (TDT) [32] or Pattern Recognition for Neuroimaging Toolbox (PRoNTo) [33], particularly designed for fMRI studies have been developed in the past years. Despite the good spatial resolution of the fMRI, the poor temporal resolution of the BOLD signal limits an accurate study of how cognitive processes unfold in time. For that reason, the application of multivariate pattern analyses to other neuroimaging techniques with a higher temporal resolution, such as EEG or magnetoelectroencephalography (MEG), is growing in popularity. With the aim of facilitating the work of researchers from different disciplines, allowing the access to these complex computation algorithms, diverse M/EEG-focused toolboxes have been developed. The Amsterdam Decoding and Modeling Toolbox (ADAM) [34], CoSMoMVPA [35], MVPA-light [36], The Decision Decoding Toolbox (DDTBOX) [37], BCILAB [38] and The Berlin Brain-Computer Interface [39] are excellent examples of MATALB-based toolboxes. MNE-Python[40], Nilearn[41] or PyMVPA [42,43] are other Python-based and open source alternatives.

### 1.2 MVPAlab: an easy-to-use machine learning toolbox for decoding analysis

Despite the tremendous effort applied in other implementations to facilitate researchers the use of these tools (e.g. high-level functions which compute a complete decoding analysis in a few lines of code), its use is sometimes really challenging, especially for students, newcomers or other researchers with profiles with no coding experience.

Here we present MVPAlab, an easy-to-use decoding toolbox for M/EEG data. So, what makes MVPAlab different from any other existing alternatives? The MVPAlab Toolbox has been designed to include an easy-to-use and very intuitive Graphic User Interface (GUI) for the creation, configuration, and execution of different decoding analysis. Importantly, this friendly GUI provides access to an extensive set of computational resources to design, configure and execute the complete pipeline of different decoding analyses for multidimensional M/EEG data, including visualization software for data representation. MVPAlab implements several decoding functionalities, such as time-resolved binary classification, temporal generalization, multivariate cross-classification, statistical analyses to find significant clusters, feature contribution analyses, and many others. Highly configurable linear and non-linear ML models can be selected as classification algorithms, including Support Vector Machines (SVM) or Discriminant Analysis (DA). Additionally, MVPAlab offers several data preprocessing routines: trial averaging, data smoothing and normalization, dimensionality reduction, among others. This MVPAlab GUI also includes a very flexible data representation utility, which generates really appealing and colorful plots and animations. In addition to this, MVPAlab implements some exclusive analyses and functionalities, such as parallel computation, which divides the computational load in different execution threads, significantly reducing the computation time, or frequency contribution analysis, which allows to estimate how relevant information is distributed across different frequency bands.

Hence, MVPAlab has not been designed for beginners only, as implements several high and low-level routines allowing more experienced profiles to design their own projects in a highly flexible manner. The following sections depict, in as much detail and as descriptively as possible, the main aspects of MVPAlab, including installation, compatibility, data structure, and a complete getting started section.

### 1.3 Installation, compatibility and requirements

The installation of MVPAlab Toolbox is quite simple. First, an up-to-date version of the code is freely available for download in the following GitHub repository:

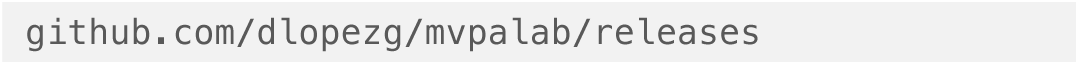

Users should (1) select and download the source code of the desired release, (2) unzip the downloaded source code folder and (3) add it to the MATLAB path with subfolders. Please see MVPAlab wiki for more detailed instructions:

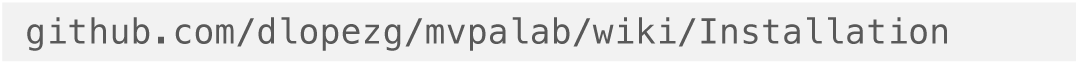

The MVPAlab Toolbox has been designed to be fully compatible with MATLAB 9.0 (R2016a) and above. This restriction is only applicable to the graphic user interface, which has been developed using App Designer, introduced in the 9.0 version. Custom MVPAlab scripts can be executed under older MATLAB versions. Other toolboxes include several function names overlapping the MATLAB (or other external packages) built-in functions, causing in some cases errors and malfunctioning. To avoid this type of problems, MVPAlab uses a specific suffix in their function names. Since this software has been developed using MATLAB and has no external dependencies, the MVPAlab Toolbox is fully supported by GNU/Linux, Unix, Windows and macOS platforms. Hardware requirements depend on the size of the analyzed dataset. While the CPU specifications only affects to the computation time, enough RAM capacity is required to store and process M/EEG data. For almost any process, the recommended RAM capacity is at least the double of the size of the dataset (measured in gigabytes). For more memory demanding processes, such as frequency contribution analysis, MVPAlab splits and stores EEG data on the hard drive, importing it again when needed. Since MVPAlab only uses the CPU for computation, the GPU specification does not affect to the toolbox performance.

Some MATLAB built-in packages and functions are required for a correct functioning of this software. For the statistical analysis, the Image Processing Toolbox is required to find clusters in significant masks. The Statistics and Machine Learning Toolbox provides functions to train and validate classification models, dimensionality reduction, feature selection, etc. The Signal Processing Toolbox is required for extracting M/EEG envelopes as features. The Parallel Computation Toolbox is not required but recommended to drastically reduce the computation time as it allows to divide the computational load in different processing threads. Finally, MVPAlab greatly benefits from other open source M/EEG toolboxes such as EEGlab and FieldTrip: some filtering functions require the EEGlab Toolbox installed and initiated for a correct operation. If MVPAlab finds an EEGlab installation it will initiate it automatically. Because of all of this, users should ensure that these dependencies are included in their MATLAB installation.

### 1.4 Dataset structure and format

MVPAlab is not a preprocessing tool for M/EEG data, instead, it is designed to read and work with epoched data from two of the most employed preprocessing toolboxes: EEGLAB and FieldTrip. For a correct operation of MVPAlab Toolbox, epoched data should be previously saved in one independent file for each subject using a `.mat` format. EEGlab format `.set` is also supported. The data structure and format should remain unaltered. If EEGlab was used for the data preprocessing, users should save the entire `EEG` structure for each participant, not only the `EEG.data` matrix. MVPAlab collects additional information from the data file, such as sampling frequency (`EEG.srate`), the location of the electrodes (`EEG.chanlocs`) or data time points (`EEG.times`). In the same way, if FieldTrip is used, users must save the entire data structure, as MVPAlab reads the required subject’s data from `data.trial`, `data.time` and `data.fsample`.

### 1.5 MVPAlab Toolbox architecture

The complete architecture of MVPAlab Toolbox is shown in Figure 1, including several of the configuration parameters, processes and routines employed for a complete decoding analysis. The complete architecture and its configuration parameters are resumed in the following stages:

**Figure 1.**
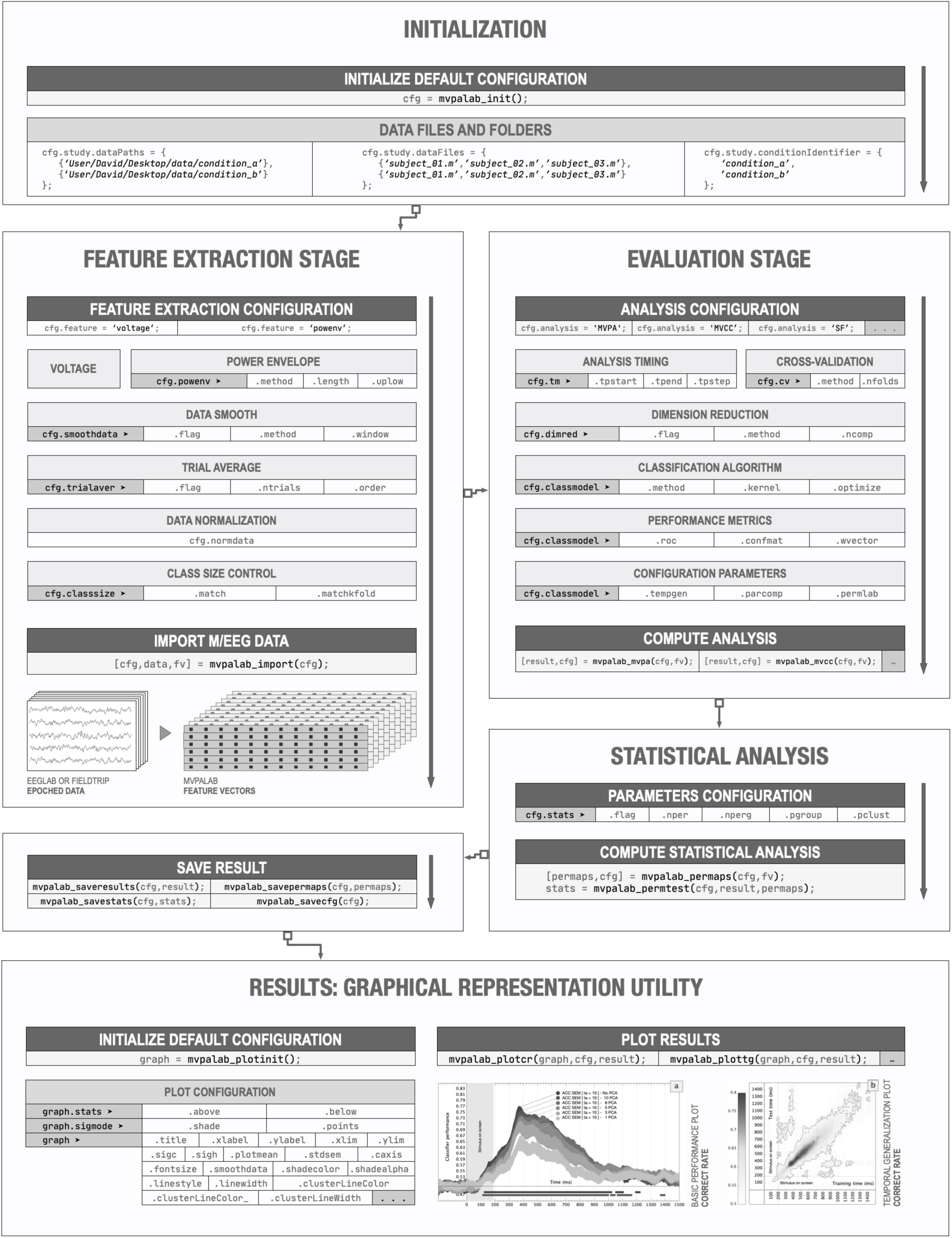
MVPAlab Toolbox complete architecture and configuration parameters.

#### Initialization stage

During the initialization stage, MVPAlab generates a default configuration structure. This variable is required for a correct operation of the toolbox.

#### Import data and feature extraction stage

Here, M/EEG data is imported, preprocessed, and prepared for the decoding analysis. During this stage, some specific configuration is required: the participants’ files to import, identifiers for binary classes, the complete path to the dataset, and others. Additionally, users can select which M/EEG feature will be extracted for classification (raw signal voltage or its envelope); enable or disable and configure several preprocessing procedures, such as trial averaging, data normalization, balanced class sizes, and others. All these preprocessing procedures are computed during this stage. Finally, the feature vectors are extracted and prepared for the multivariate analysis.

#### Evaluation stage

During the evaluation stage, several classification models can be trained and validated using cross-validation approaches. Dimensionality reduction, if enabled, is also computed during this stage.

Users can specify different classification models, linear and non-linear kernel functions, different cross-validation techniques, different model’s performance metrics, etc. The results of the decoding analysis, the configuration file and other analysis-related files will be hierarchically stored in the project’s directory. This directory is the folder containing the main analysis script.

#### Statistical significance stage

If permutation test is enabled, statistically significant clusters are extracted from the result via non-parametric cluster-based permutation testing. For this stage, users can specify the total number of permutations at a participant and group level to be computed, the p-value thresholds for a data point or cluster size to be considered significant and other relevant information.

#### Graphical representation stage

Last but not least is the graphical representation stage. MVPAlab has fully integrated high-level plotting tools, allowing researchers to easily design and generate high quality and highly customizable result representations.

### 1.6 Getting started

Computing a multivariate analysis in MVPAlab Toolbox is quite simple for all type of users. Researchers with no coding experience can use the integrated graphic user interface, which allows to create, save, configure, execute and plot the results of any supported multivariate analysis in a very intuitive way. Not a single line of code is needed. However, users with coding experience looking for a faster and more flexible way to interact with the toolbox can create their own scripts. Be that as it may, MVPAlab also includes several easy-to-understand and well-documented demo scripts for different types of analyses, making this tool very convenient not just for experienced users but also for newcomers. This section includes a general introduction to the functioning of MVPAlab Toolbox, either by using the GUI or building custom scripts.

#### 1.6.1 GRAPHIC USER INTERFACE

Once MVPAlab is installed, the graphic user interface can be launched by typing the following command in the MATLAB command line:

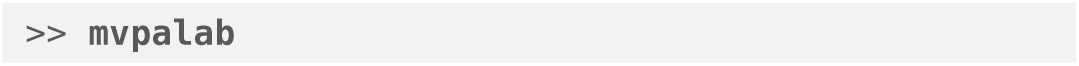

##### Creating new analyses

If the MVPAlab folder is correctly added to the MATLAB path as described in Section 1.3, the initial MVPAlab window should appear as shown in Figure 2 (a). Using this interface, users can create new analyses, open previously created analyses or open the plotting utility. Creating new analyses in MVPAlab using the GUI is very simple and intuitive. Researchers only need to specify the type of analysis required from the dropdown menu and select the location folder. Results, configuration and other analysis-related files will be hierarchically stored in this directory. Once everything is selected, clicking the configuration button will create the project folder structure and launch the analysis configuration window, as shown in Figure 2 (b).

**Figure 2.**
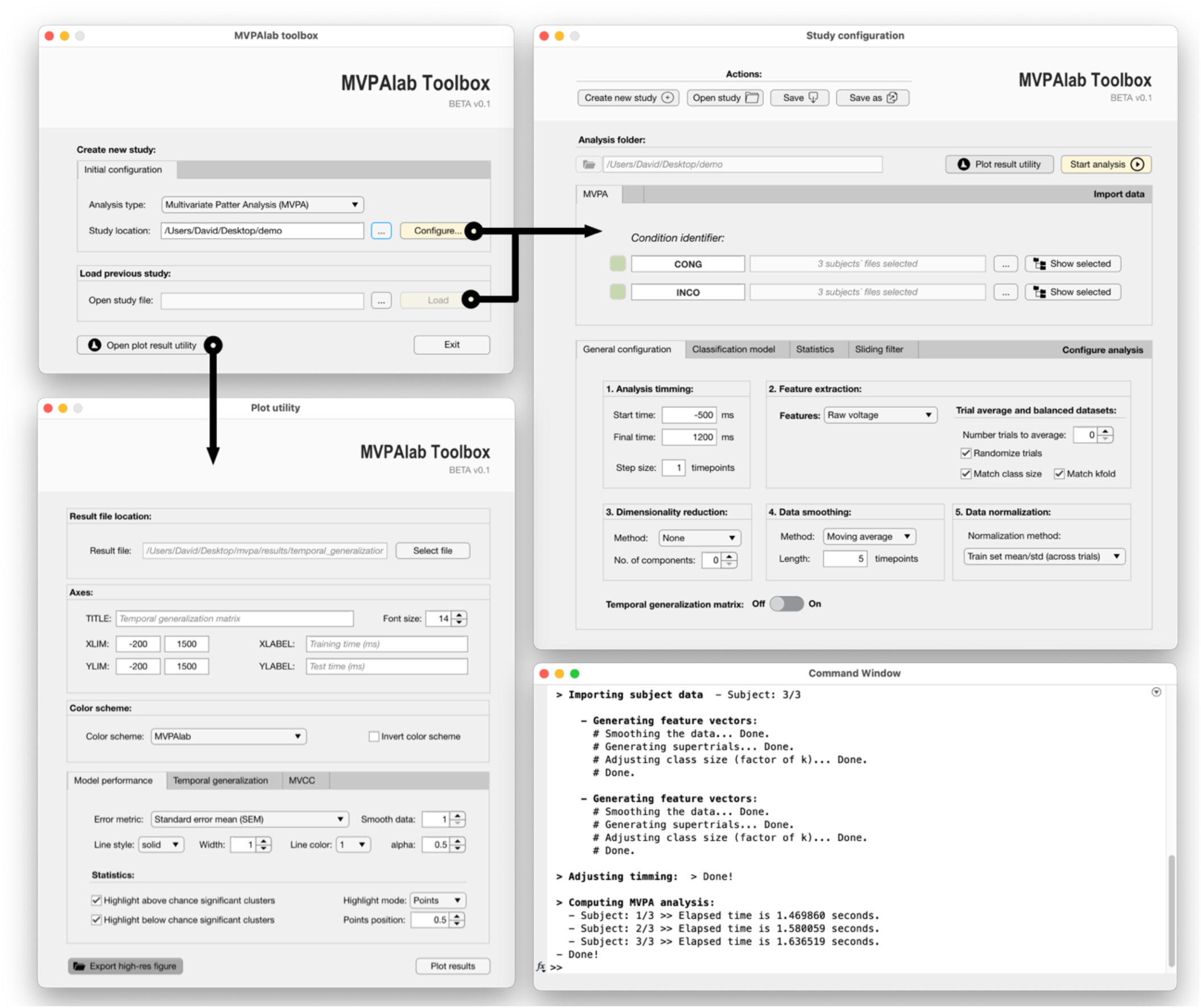
MVPAlab graphic user interface. (a) Initial view. (b) Analysis configuration view. (c) Plot utility view. (d) MATLAB command window.

##### Configuring the decoding analysis

Before computing a multivariate analysis, additional details of configuration are required. Users must specify the locations of the epoched datasets and label each condition with a condition identifier. All the relevant parameters of the decoding analysis are set to its default value and can be modified within this configuration window. These configuration parameters include a wide range of processes that can be executed during the decoding analysis, such as: data normalization, data smoothing, trial averaging, analysis timing, dimensionality reduction, balance datasets and others. Additionally, the employed classification models can also be designed here. Users can choose between different classification algorithms, kernel functions, cross-validation strategies and select several output performance metrics. They can enable the computation of the temporal generalization matrix, activate parallel computation or configure statistical analyses. All MVPAlab toolbox functionalities are perfectly detailed in Section 2 *Materials and Methods*.

##### Computing the decoding analysis

Once the configuration parameters are correctly specified, the computation of the multivariate analysis can be started by clicking the *Start analysis* button. Depending on the size of the dataset and the selected configuration, this process may be time-consuming and CPU/memory demanding. Anyhow, during the computation of the entire analysis pipeline, as shown in Figure 2 (c), MVPAlab prompts in the MATLAB command window detailed information of the processes being executed.

##### Plotting the results

For the graphical representation of the results, MVPAlab also offers an intuitive plot utility that can be opened by clicking on *Open plot utility* button Figure 2 (d). This tool enables users to open, plot, combine and compare results of different analyses without dealing with cumbersome lines of MATLAB code. The most common configuration parameters such as titles, labels, line styles, transparencies, color palettes, axes limits, data smoothing or highlighting can be easily configured for time-resolved analysis, temporal generalization matrices, frequency contribution analyses, and others. In addition, with this interface users can create animated temporal representations of feature weights distribution over scalp templates.

All this combined allows researchers with no or little coding experience to prepare and compute multivariate decoding analyses of M/EEG data; create high quality and ready-to-publish figures, all of this without witting a single line of code.

#### 1.6.2 BUILDING CUSTOM SCRIPS

The intuitive and easy-to-use GUI is not the only way to utilize this software. For those researchers looking for flexibility and automation, MVPAlab implements several high-level functions to easily set up a custom decoding analysis. The complete analysis pipeline can be divided into five main steps, including the statistical permutation test and plotting functions, and runs as follows:

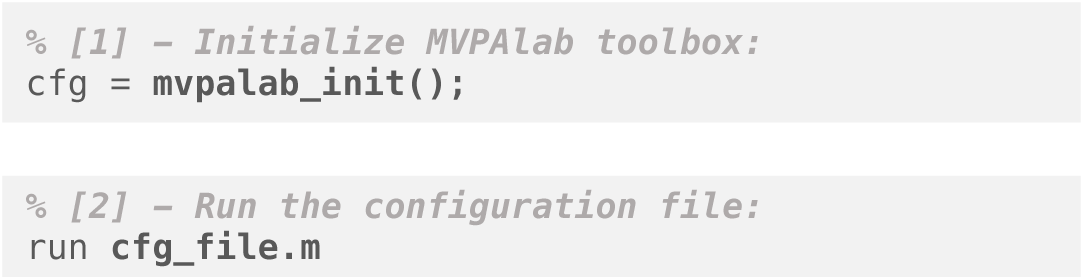
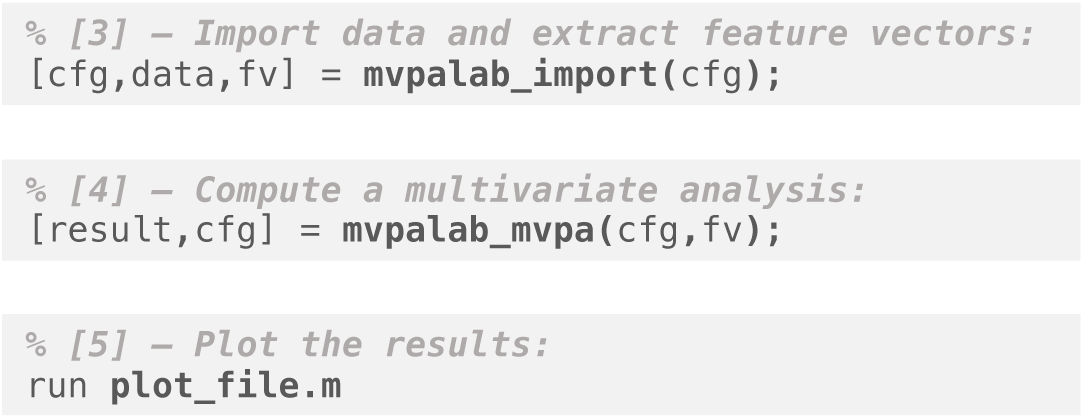

First, the function `mvpalab_init()` initializes the toolbox. This function returns a default configuration structure `cfg`, which consist of all the required configuration parameters for an analysis. Please see the *Material and Methods* section for a detailed description of each field of the configuration variable.

Users should modify this configuration variable to set up the desired configuration for a specific decoding analysis. For the sake of clarity and for maintaining a clean code organization, all this configuration code should be placed in an external configuration file `cfg_file.m`. This file will be executed after the toolbox initialization.

Once the MVPAlab toolbox is initialized and a specific analysis configured, the function **mvpalab_import(**cfg) imports and preprocess the datasets provided, according to the configuration file `cfg`. This function returns a copy of the preprocessed data (`data`), which can be omitted to save memory, and the extracted feature vectors (`fv`), which will be the input for the classification models. Please see Section 2.3 *Importing Data and Feature Extraction* for a more detailed explanation of the feature extraction process.

Next, the function **mvpalab_mvpa(**cfg,fv) computes the multivariate pattern analysis. Other functions are available for different analyses, such as **mvpalab_mvcc(**cfg,fv) for cross-classification and **mvpalab_sfilter(**cfg,fv) for frequency contribution analysis.

These functions return the variable `result`, which includes the time-resolved decoding performance for every performance metric enabled in the configuration file. In addition, the result files are automatically saved in separate folders in the project directory.

To compute the statistical analysis and draw statistical inferences at the group level, one additional step should be added to the former execution pipeline:

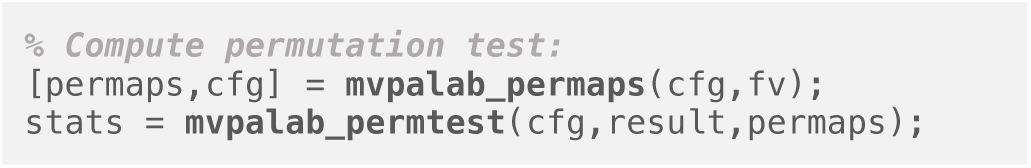

These functions implement a non-parametric cluster-based permutation test, returning the variable `stats`, which includes statistically significant clusters found in the data. Please, see Section 2.5 *Cluster-based permutation testing* for an exhaustive description of this test.

Finally, in addition to the graphic user interface, MVPAlab includes several plotting routines, allowing users to design customizable and ready-to-publish figures and animations. Please see Section 2.6 *Result representation pipeline* for more details. Several demo scripts for different types of analyses and result representations are included in the MVPAlab Toolbox folder.

## 2. MATERIALS AND METHODS

### 2.1 Sample EEG dataset

A sample EEG dataset has been compiled to test all the MVPAlab main functionalities. It is freely available in the following repository:

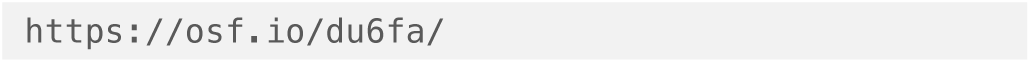

Here, three different EEG data files have been selected from the original work [44,45]. For each participant, two different main conditions (*condition_a* vs. *condition_b*) have been selected for the MVPA analysis. Additionally, four subconditions (*condition_1, condition_2*, vs. *condition_3 and condition_4*) have been selected for the multivariate cross-classification analysis. Readers interested on the experimental details of these data should refer to the original publication [44,45]. During the original study, high-density EEG was recorded from 65 electrodes. The TP9 and TP10 electrodes were used to record the electrooculogram (EOG) and were removed from the dataset after the preprocessing stage. Impedances were kept below 5kΩ and EEG recordings were average referenced, downsampled to 256 Hz, and digitally filtered using a low-pass FIR filter with a cutoff frequency of 120 Hz, preserving phase information. No channel was interpolated for any participant. Continuous data were epoched [−1000, 2000ms centered at onset of the stimulus] and baseline corrected [−200, 0ms]. Independent Component Analysis (ICA) was computed to remove eye blinks from the signal, and the artifactual components were rejected by visual inspection of raw activity of each component, scalp maps and power spectrum. Finally, an automatic trial rejection process was performed, pruning the data from non-stereotypical artifacts. For more details please see [44].

The final compiled dataset consists of an EEGlab data structure per subject and condition with **[63** x **768** x **ntrials]** EEG data matrices. The number of trials per condition and participant is shown in the following table:

**Table 1.** Total number of trials per subject and condition.

|  | subject_01.mat | subject_02.mat | subject_03.mat |
| --- | --- | --- | --- |
| condition_a | 468 | 413 | 434 |
| condition_b | 403 | 399 | 396 |
| condition_1 | 212 | 193 | 190 |
| condition_2 | 218 | 202 | 212 |
| condition_3 | 191 | 206 | 206 |
| condition_4 | 250 | 211 | 222 |

### 2.2 Defining a configuration file

For the sake of clarity and code organization, we recommend to include all the configuration code for a specific decoding analysis in an external configuration `.m` file. This file should be executed before the computation of the multivariate decoding analysis. This recommendation, however, is not mandatory and more experienced users can design their own scripts according to their needs and preferences. For both scenarios, all the available configuration parameters in MVPAlab Toolbox will be described in detail during this section.

#### 2.2.1 PARTICIPANTS AND DATA DIRECTORIES

The first required information that should be specified by the user is the working directory and the location of the dataset to be imported and analyzed. This includes, for each class or condition, the name of each individual subject data file and the complete path of the class folder. These parameters can be defined in the configuration file as follows:

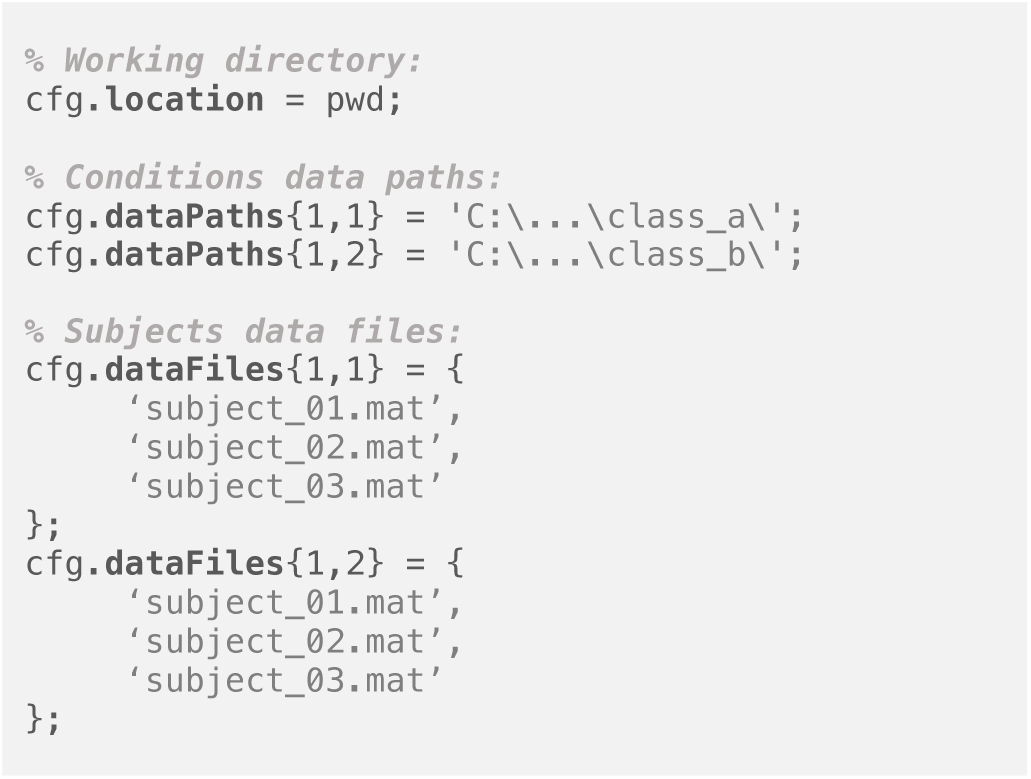

Before computing the multivariate decoding analysis, the MVPAlab Toolbox can be used to execute several preprocessing procedures that may improve the final results in different ways (e.g. increasing accuracy, avoiding skewed results, data normalization, data smoothing, etc.). The default configuration of each of these procedures is initialized when MVPAlab toolbox is launched. However, these procedures and their configuration parameters can be adjusted by the users to meet the required specific analysis conditions. During this section, all of these preprocessing procedures and their configuration parameters will be meticulously described.

#### 2.2.2 TRIAL AVERAGING

If enabled, this approach randomly or sequentially averages a certain number of trials **n**_trials_ belonging to the same condition for each participant. This procedure creates *supertrials* and usually increases the signal-to-noise ratio (SNR) which improves the overall decoding performance and reduces the computational load. Since reducing the number of trials per condition typically increases the variance in the decoding performance, this procedure imposes a trade-off between the increased variance/accuracy. It should be noted that increasing **n**_**trials**_ does not increase the decoding performance linearly. Please see [46,47] for more details.

The default parameters for this procedure can be modified in the MVPAlab configuration file as follows:

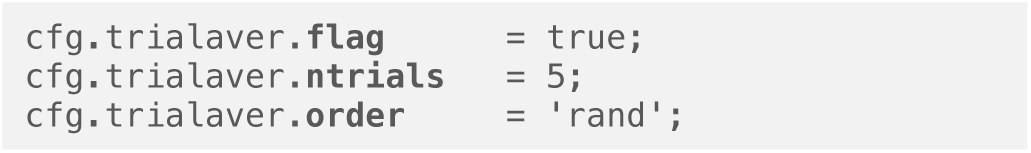

Trial averaging can be enabled or disabled by setting the configuration variable (.**flag)** to true or false. The number of trials to average can be modified in (.**ntrials)**. Finally, the order in which the trials are selected for averaging can be modified setting the variable (.**order)** to ‘rand’ or ‘sequential’.

#### 2.2.3 BALANCED DATASETS

Unbalanced datasets can lead to skewed classification results [48]. To avoid this phenomenon, the number of trials per condition should be the same. MVPAlab can be used to define strictly balanced datasets by downsampling the majority class to match the size of the minority one (cfg.classsize.**match**). In addition, each class size can be set as a factor of k, the total number of folds in the cross-validation (CV) procedure. Thus, during CV each fold will be composed by exactly the same number of observations, avoiding any kind of bias in the results (cfg.classsize.**matchkfold**).

These features are disabled by default but can be enabled in the MVPAlab configuration structure as follows:

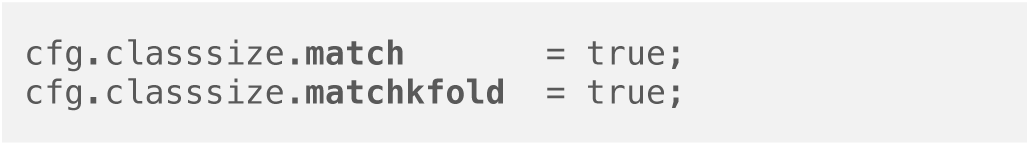

#### 2.2.4 DATA NORMALIZATION

In machine learning, data normalization refers to the process of adjusting the range of the M/EEG raw data to a common scale without distorting differences in the ranges of values. Although classification algorithms work with raw values, normalization usually improves the efficiency and the performance of the classifiers [49]. Four different (and excluding) data normalization methods are implemented in MVPAlab. A commonly used normalization approach [50] is computed within the cross-validation loop. Hence, the training and test sets are standardized as follows:

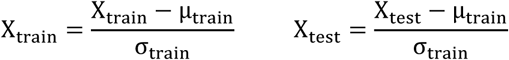

where μ_train_ and σ_train_ denote the mean and the standard deviation of each feature (column) of the training set. Other normalization methods implemented in MVPAlab are: z-score (μ = 0 ; σ = 1) across time, trial or features. To compute these normalization strategies MVPAlab uses the MATLAB built-in function *zscore*, included in the Statistics and Machine Learning Toolbox.

Data normalization method, which is disabled by default, can be modified as follows:

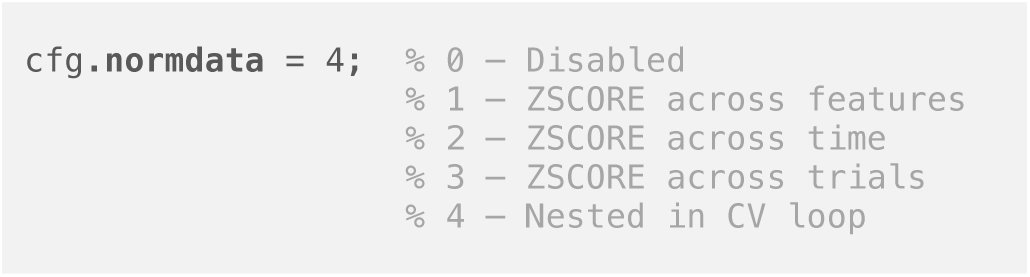

#### 2.2.5 DATA SMOOTHING

Data smoothing is a procedure employed in recent M/EEG studies [51–54] to attenuate unwanted noise. MVPAlab implements an optional data smoothing step that can be computed before multivariate analyses. This procedure is based on MATLAB built-in function *smooth*, included in the Curve Fitting Toolbox, which smooths M/EEG data points using a moving average filter.

The length of the smoothing window can be specified in the variable (cfg.smoothdata.**window**) and should be an odd number. For a window length of 5 time points, the smoothed version of the original signal is computed as follows:

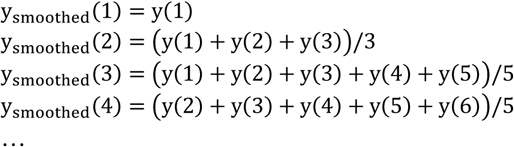

Data smoothing is disabled (.**method** = ‘none’) by default and can be enabled and configured in the MVPAlab configuration file as follows:

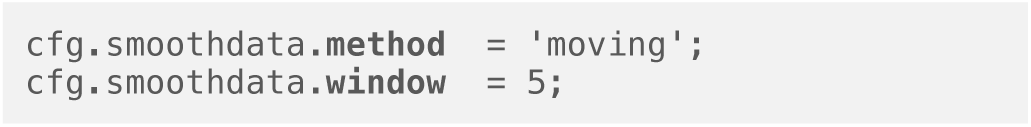

#### 2.2.6 ANALYSIS TIMING

By default, MVPAlab computes the time-resolved decoding analysis for each timepoint across the entire M/EEG epoch. However, the user can define a specific region of interest (time window) and a different step size as follows:

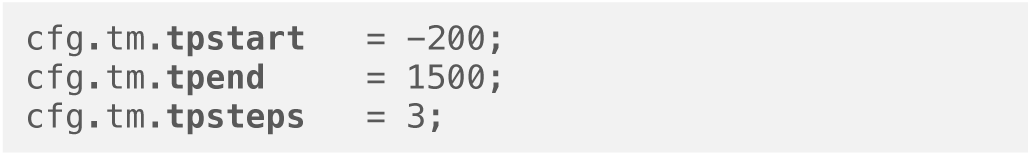

This way, the temporal decoding analysis will be computed from -200ms (.**tpstart**) to 1500ms (.**tpend**) not for each timepoint but for every three (.**tpsteps**) timepoints. Note that increasing the step size decreases the processing time but also causes a reduction in the temporal resolution of the decoding results.

#### 2.2.7 DIMENSIONALITY REDUCTION

In machine learning, dimension reduction techniques are a common practice to reduce the number of variables in high-dimensional datasets. During this process, the features contributing more significantly to the variance of the original dataset are automatically selected. In other words, most of the information contained in the original dataset can be represented using only the most discriminative features. As a result, dimensionality reduction facilitates, among others, classification, visualization, and compression of high-dimensional data [55]. There are different dimensionality reduction approaches but Principal Component Analysis (PCA) is probably the most popular multivariate statistical technique used in almost all scientific disciplines [56], including neuroscience [57].

PCA in particular is a linear transformation of the original dataset in an orthogonal coordinate system in which axis coordinates (principal components) correspond to the directions of highest variance sorted by importance. To compute this transformation, each row vector **x**_**i**_ of the original dataset **X** is mapped to a new vector of principal components **t**_**i**_ = (**t**_**1**_, …, **t**_**1**_), also called scores, using a p-dimensional coefficient vector **w**_**j**_ = (**w**_**1**_, …, **w**_**p**_):

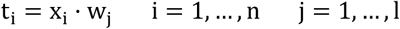

For dimension reduction: **1** < **p**.

To maintain the model’s performance as fair and unbiased as possible, PCA is computed only for training sets **X**_**training**_, independently for each fold inside the cross-validation procedure. Once PCA for the corresponding training set is computed and the model is trained, the exact same transformation is applied to the test set **X**_**test**_ (including centering, **μ**_**training**_). In other words, the test set is projected onto the reduced feature space obtained during the training stage. According to the former equation, this projection is computed as follows:

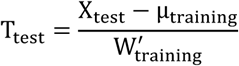

To compute this nested implementation of the PCA algorithm, MVPAlab uses the MATLAB built-in function *pca*, included in the Statistics and Machine Learning Toolbox. However, dimensionality reduction techniques such PCA endorse a tradeoff between the benefits of dimension reduction (reduced training time, reduced redundant data and improved accuracy) and the interpretation of the results when electrodes are used as features. When PCA is computed, the data is projected from the sensor space onto the reduced PCA features space. This linear transformation implies an intrinsic loss of spatial information, which means that, for example, we cannot directly analyze which electrodes are contributing more to decoding performance. The default parameters for this procedure can be modified in the MVPAlab configuration file as follows:

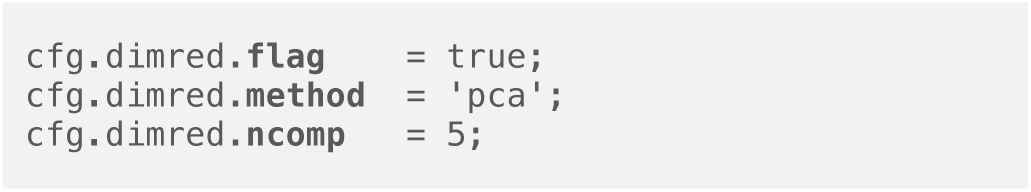

#### 2.2.8 CLASSIFICATION ALGORITHMS

Classification algorithms are the cornerstone of decoding analyses. These mathematical models play the central role in multivariate analyses: detect subtle changes in patterns in the data that are usually not detected using less sensitive approaches. Different classification algorithms have been used to achieve this goal, from probabilistic-based models such as Discriminant Analyses (DA), Logistic Regressions (LR) or Naïve Bayes (NB) to supervised learning algorithms such Support Vector Machine (SVM).

For the time being, MVPAlab Toolbox implements two of the most commonly employed models in the neuroscience literature, Support Vector Machines and Discriminant Analysis in their linear and non-linear variants.

The classification model employed for the decoding analysis can be specified in the configuration file as follows:

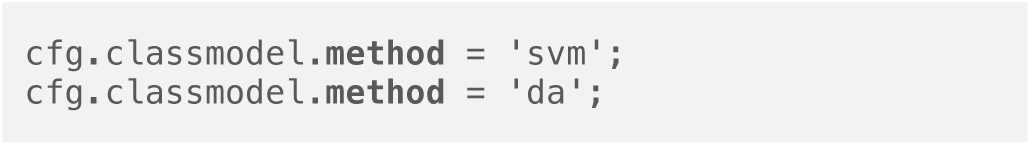

Both classification approaches are based on MATLAB built-in libraries for support vector machines and discriminant analyses. A brief mathematical description for both models can be found below. Please see the MATLAB documentation of `fitcsvm` and `fitcdiscr` functions for further details.

##### Support Vector Machine

Support Vector Machine (SVM) provides a theoretically elegant, computationally efficient, and very effective solution for many practical pattern recognition problems [58–60]. For that reason, SVM is broadly employed in M/EEG studies. Intuitively, for binary classification problems, during the training stage this algorithm searches for an optimal hyperplane maximizing the separation between this hyperplane and the closest data points of each class. These data points are called *support vectors*. The separation space is called *margin* and is defined as 2/‖**w**‖, and it does not contain any observation for separable classes, as shown in Figure 3 (a). Thus, the linear SVM score function is defined as follows:

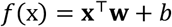

where the input vector **x** is an observation, the vector **w** contains the coefficients that define an orthogonal vector to the hyperplane and *b* is the bias term. To formalize the optimization problem (that is, to find the optimal hyperplane that maximizes the margin), several constraints should be defined. Therefore, any given sample will be correctly classified as long as:

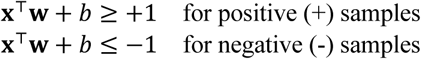

**Figure 3.**
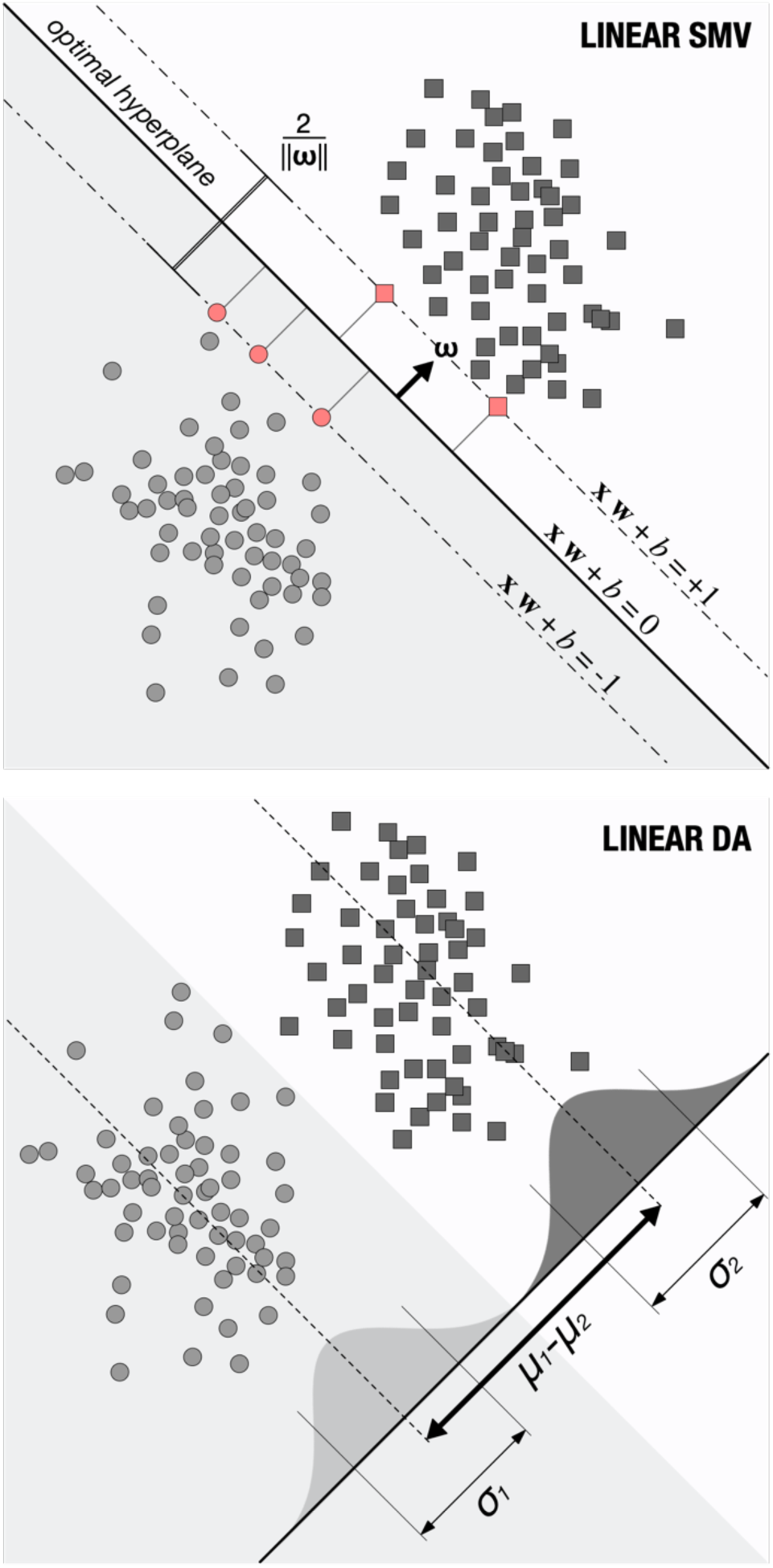
Classification models: graphical representation of (a) LSVM and (b) LDA classifiers for simulated data. Red points represent the support vectors, the closest data points to the decision boundary (hyperplane).

Introducing y_1_ = {+1, −1} for positive and negative samples, respectively, the two former equations can be rewritten for mathematical convenience as follows:

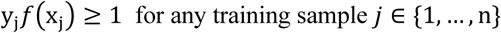

This is the decision rule for separable classes. When the classes are not perfectly separable, the algorithm imposes a penalty introducing positive slack variables ξ_1_ > 0 for each observation on the wrong side of the hyperplane. For those observations that are correctly placed: ξ_1_ = 0. Consequently, non-separable data impose a trade-off between margin maximization and the total number of constraint violations. Now, the optimization problem reads as follows:

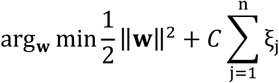

with respect to **w** and *b* and subject to:

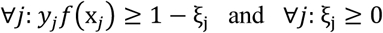

The parameter *C* is a constant which modulates the trade-off between the training error and the complexity of the model. A search-grid-based optimization of the misclassification cost parameter *C* can be enabled and computed using five-fold CV for the training set on the configuration file as follows:

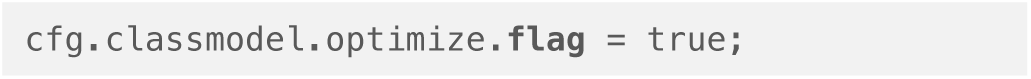

For some classification scenarios, it is not always possible to find an optimal criterion for class separation using linear classifiers. To solve this problem, original data from the input space 𝒩 can be mapped into a high dimensional feature space ℱ using a mapping function *ϕ*: 𝒩 ⟼ ℱ. Therefore, the decision equation is now defined as follows:

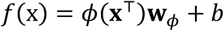

However, the application of the transformation function *ϕ* is not explicitly needed. Since the hyperplane optimization problem depends on nothing but pairwise dot products (e.g. **x**_1_ · **x**_2_), we only need a set of kernel functions that meet the following property: *K*(**x**_1_, **x**_2_) = ⟨*ϕ*(**x**_1_), *ϕ*(**x**_2_)⟩.

This class of function includes, among others, polynomial or gaussian kernels:

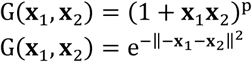

The mentioned variant of the initial mathematical approach for non-linear classifiers is known as *kernel trick* (Figure 4) and it retains nearly all the simplicity and benefits of linear approaches, making data linearly separable in the feature space ℱ. However, in decoding analyses, linear approaches are normally preferred not just for their simplicity, but also for yielding comparable performance results in several applications [61].

**Figure 4.**
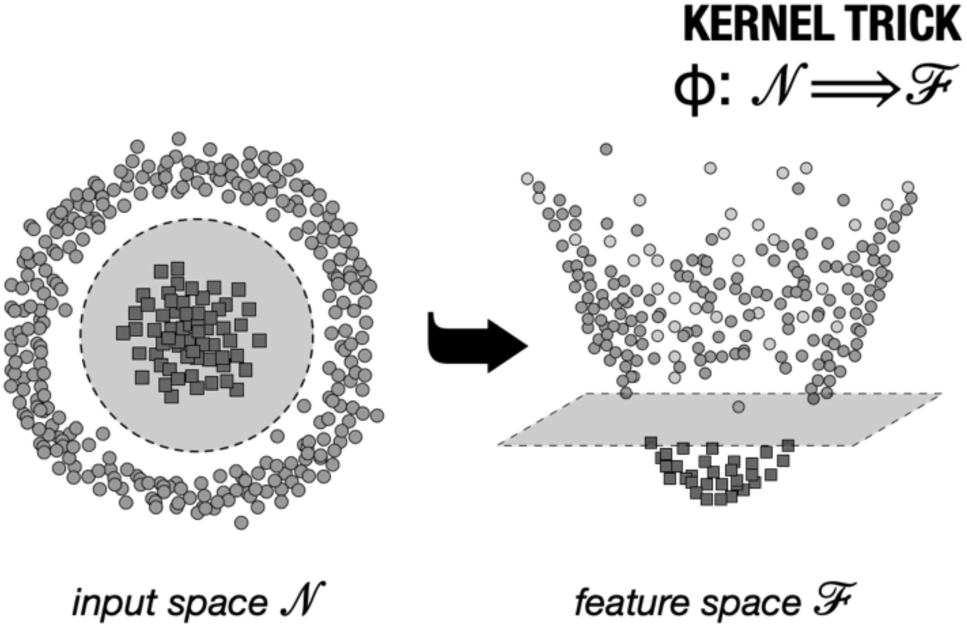
Kernel trick graphical representation: original data in the input space is not linearly separable. This data points can be projected into a high-dimensional space using the mapping function ϕ. In this new feature space, classes became separable using linear approaches.

MVPAlab uses linear classifiers for decoding analysis by default, but other kernel functions for non-linear classification can be specified in the MVPAlab configuration file as follows:

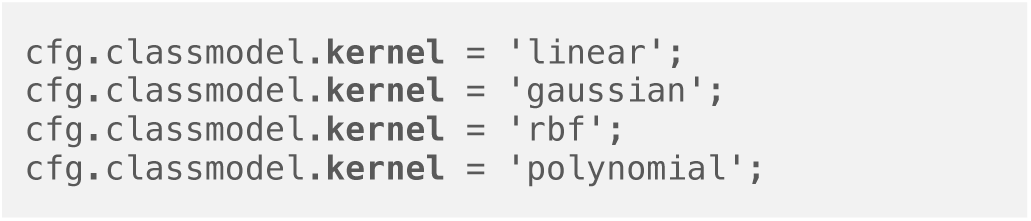

##### Discriminant analysis

Prediction using Discriminant Analysis (DA), see Figure 3 (b), is based in three different metrics: posterior probability, prior probability and cost. Thus, the classification procedure tries to minimize the expected classification cost:

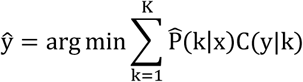

where ŷ is the predicted classification, **K** corresponds to the number of classes, 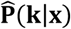 is the posterior probability of class **k** for observation **x** and C(y|k) is the cost of classifying an observation as **y** when its true class is **k**.

Being **P**(**k**) the prior probability of class **k**, the posterior probability that an observation **x** belongs to class **k** is:

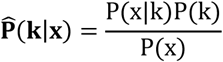

where:

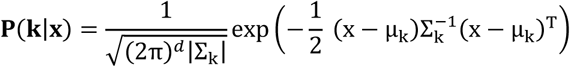

is the multivariate normal density function, being **Σ**_**k**_ the ***d***-by-***d*** covariance matrix and **μ**_**k**_ the 1-by-***d*** mean. Please see the MATLAB documentation for further details.

While Linear Discriminant Analyses (LDA) assumes that both classes have the same covariance matrices **Σ**_**k**_ and only the means **μ**_**k**_ vary, for Quadratic Discriminant analyses (QDA), both means and covariance matrices may vary. Thus, decision boundaries are determined by straight lines in LDA and by conic sections (ellipses, hyperbolas or parabolas) for QDA. Linear Discriminant analysis is configured by default in MVPAlab Toolbox but, as for SVM, this kernel function can be modified in the configuration file as follows:

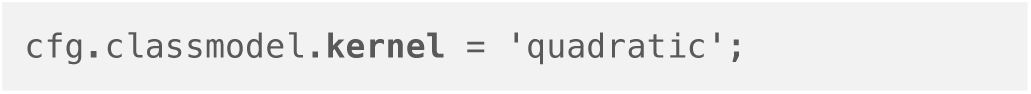

#### 2.2.9 CROSS-VALIDATION

In prediction models, cross-validation techniques are used to estimate how well the classification algorithm generalizes to unknow data. Two popular approaches for evaluating the performance of a classification model on a specific data set are *k-fold* and *leave-one-out* cross validation [62]. In general, these techniques randomly split the original dataset into two different subsets, the training set **X**_**training**_**: 1** − **1**/**K** percent of the exemplars, and the test set **X**_**test**_ : **1**/**K** percent of the exemplars. This procedure is repeated **K** times (folds), selecting different and disjoint subsets for each iteration. Thus, for each fold, the classification model is trained for the training set and evaluated using exemplars belonging to the test set. The final classification performance value for a single timepoint is the mean performance value for all iterations.

When **K** and the total number of exemplars (instances) are equal, this procedure is called *leave-one-out* cross-validation. Here, the classification model is trained with all but one of the exemplars and evaluated with the remaining exemplar. By definition, this approach is computationally demanding and time consuming for large datasets, and for that reason it is usually employed only with small sets of data. Additionally, the leave-one-out procedure has been proved to yield unstable and biased results, which makes random splits methods the preferred alternative [63].

The cross-validation procedure can be tuned in the MVPAlab configuration file as follows:

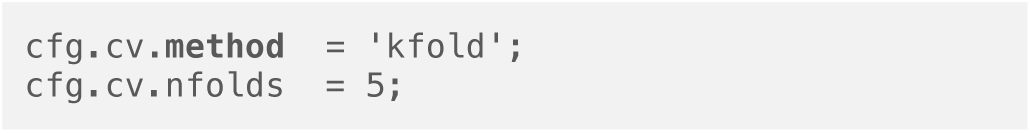

If (.**method** = ‘loo’) the number of folds is automatically updated to match the total number of exemplars for each participant

#### 2.2.10 PERFORMANCE METRICS

*(1) Mean accuracy* is usually employed to evaluate decoding models’ performance in neuroscience studies [64]. This metric is fast, easy to compute and is defined as the number of hits over the total number of evaluated trials. By default, MVPAlab Toolbox returns the mean accuracy value as a measure of decoding performance. Nevertheless, in situations with very skewed sample distributions, this metric may generate systematic and undesired biases in the results. Other performance metrics, such as the balanced accuracy have been proposed to mitigate this problem [65].

Accuracy values can be complemented with the *(2) confusion matrices*, which are very useful for binary classification but even more so for multiclass scenarios. In machine learning, a confusion matrix allows the visualization of the performance of an algorithm (see Figure 5), reporting false positives (FP), false negatives (FN), true positives (TP), and true negatives (TN). To this end, a confusion matrix reflects the predicted versus the actual classes. Rows correspond to true class and columns to predicted classes. Thus, the element **CM**_**i**,**j**_ indicates the number (or the proportion) of exemplars of class **i** classified as class **j**. Other interesting and more informative performance metrics available in MVPAlab are derivations of the confusion matrix:

(3) *Precision* **PR** = **TP**/(**TP** + **FP**): proportion of trials labeled as positive that actually belong to the positive class.

(4) *Recall* (also known as *sensitivity*) **R** = **TP**/(**TP** + **FN**): proportion of positive trials that are retrieved by the classifier.

(5) *F1-score* **F1** = **2TP**/(**2TP** + **FP** + **FN**): combination of precision and recall in a single score through the harmonic mean.

**Figure 5.**
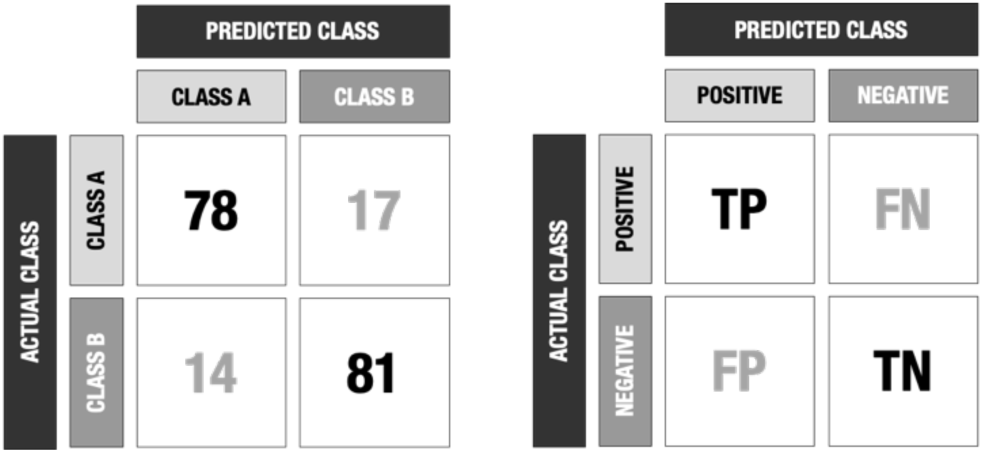
Confusion matrix. Example of a confusion matrix returned by MVPAlab Toolbox for a binary classification scenario.

Nonetheless, nonparametric, criterion-free estimates, such as the Area Under the ROC Curve (AUC), have been proved as a better measure of generalization for imbalanced datasets [66]. This curve is used for a more rigorous examination of a model’s performance. The AUC provides a way to evaluate the performance of a classification model: the larger the area, the more accurate the classification model is. This metric is one of the most suitable evaluation criteria, as it shows how well the model distinguishes between conditions, by facing the sensitivity (True Positive Rate (TPR)) against 1-specificity (False Positive Rate (FPR)), defined as follows:

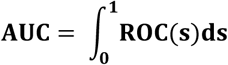

To compute the AUC and the ROC curve MVPAlab utilizes the MATLAB built-in function *perfcurve*, included in the Statistics and Machine Learning Toolbox.

By default, MVPAlab only returns the mean accuracy, although other performance metrics can be enabled in the configuration file as follows:

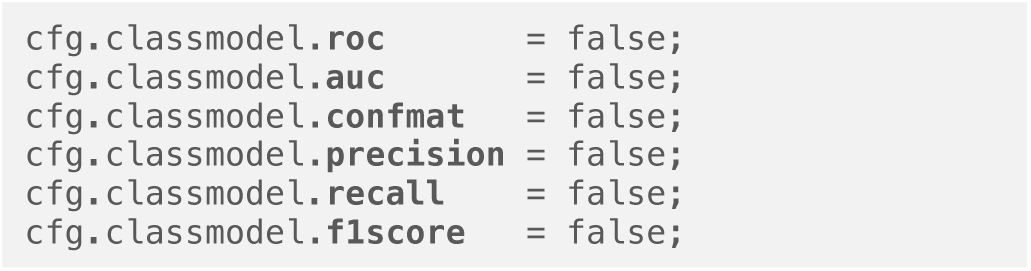

Users should be aware that enabling several performance metrics will significantly increase the computation time and memory requirements to store the results.

#### 2.2.11 PARALLEL COMPUTATION

The MVPAlab Toolbox is adapted and optimized for parallel computation. If the Parallel Computing Toolbox (MATLAB) is installed and available, MVPAlab can compute several timepoints simultaneously. Therefore, the computational load is distributed among the different CPU cores, significantly decreasing the processing time. This feature becomes critical specially when the user is dealing with large datasets and needs to compute several thousand of permutation-based analyses. Parallel computation is disabled by default but can be enabled in the MVPAlab configuration file as follows:

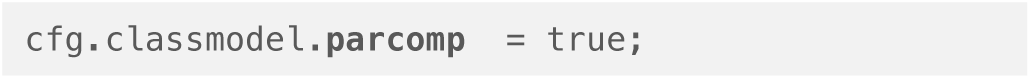

### 2.3 Importing data and feature extraction

To obtain the classification performance in a time-resolved way, the epoched M/EEG data must be prepared for the classification process. During the feature extraction step, feature vectors are defined as a selection/combination of variables of the original dataset. Typical multivariate analyses use the raw voltage of the signal as a feature for the classification, but other characteristics, such the power envelope of the signal, can also be used as features. These feature vectors are extracted as shown in Figure 6. For each participant, time-point and trial, two feature vectors (one for each condition or class) are generated, consisting of the raw potential (or any other feature such the power envelope) measured in all electrodes.

**Figure 6.**
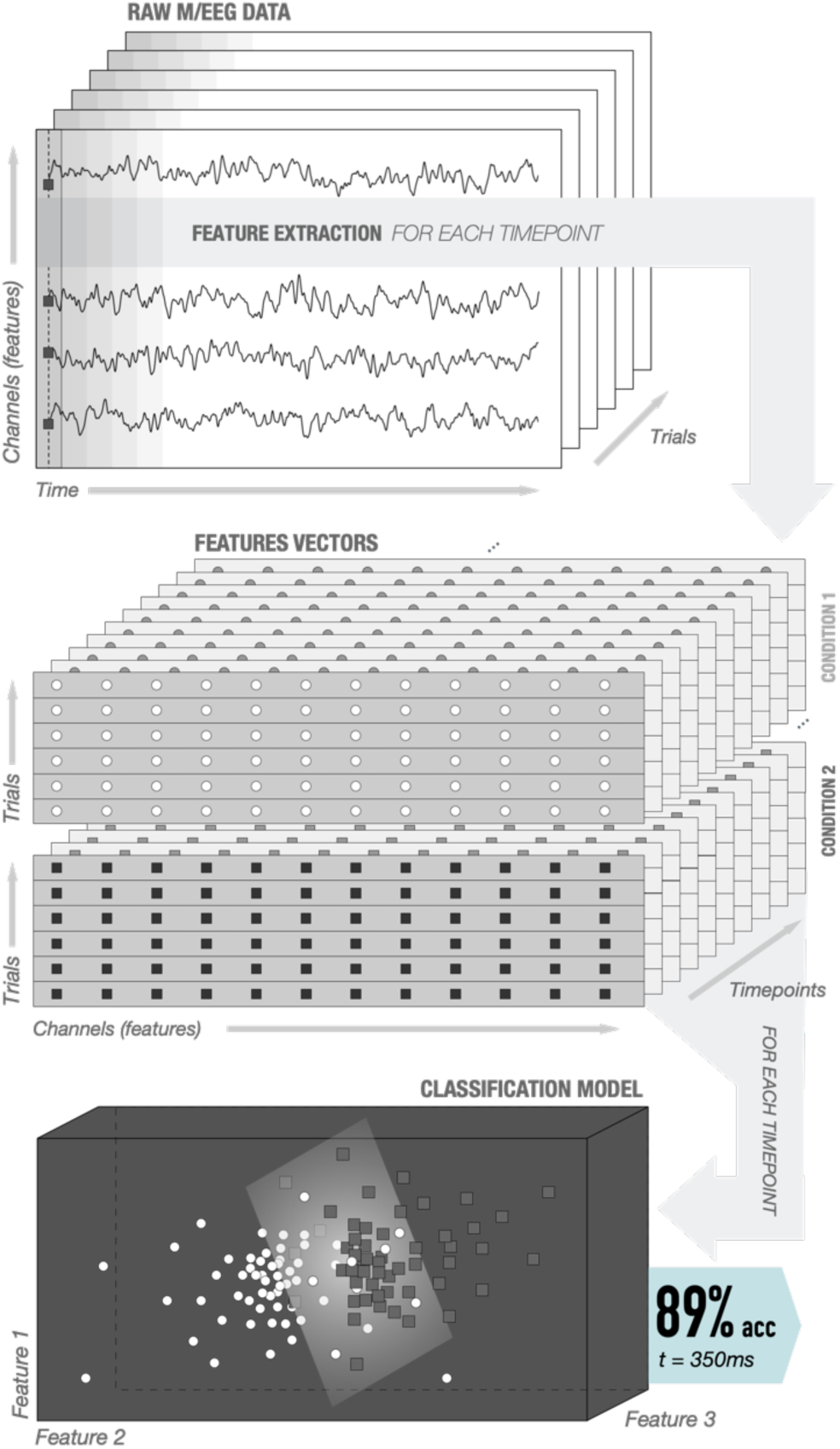
Feature extraction stage: For each participant, time-point and trial, two feature vectors are generated, one for each condition or class. These feature vectors consist of the raw potential (or any other feature such the power envelope) measured in all electrodes.

**Figure 7.**
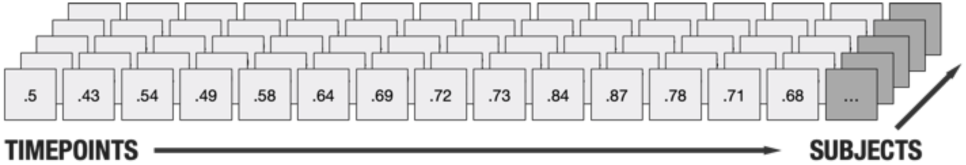
Data structure of the result file. Performance values are stored in 1 x timepoint x subject matrices. Group-level performance values can be calculated computing the mean across the third dimension.

Once MVPAlab is initialized and the analysis configuration parameters are defined in `cfg_file.m`, the function **mvpalab_import(**cfg) imports the original dataset and returns an updated version of the configuration structure (`cfg`), the preprocessed data (`data`) and feature vectors (`fv`):

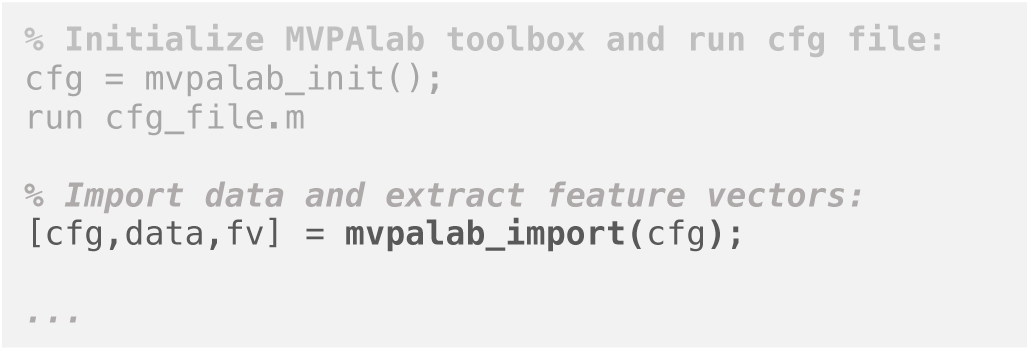

The feature vector and data variables are cell arrays structured as follows: `[1 x subjects]`. Each cell in `fv` contains a data matrix (`X`) with the feature vectors of individual subjects `[trials x features x timepoints]` and a logical vector (`Y`) including the true labels of the subject’s dataset. The data variable contains, for each condition, a data matrix including the preprocessed dataset `[features x timepoints x trials]`.

### 2.4 Type of analysis

The MVPAlab Toolbox computes two main analyses: time-resolved Multivariate Pattern Analysis (TR-MVPA) and time-resolved Multivariate Cross-Classification (TR-MVCC). Different types of analyses such the Temporal Generalization, the Feature Contribution Analysis or the Frequency Contribution Analysis are derived from them.

#### 2.4.1 TIME-RESOLVED MULTIVARIATE PATTERN ANALYSIS (TR-MVPA)

Multivariate Pattern Analyses, also known as decoding analyses, comprise a set of machine learning models that extract information patterns from multi-dimensional data. One of the most remarkable advantages of these multivariate over univariate techniques is its sensitivity in detecting subtle changes in the patterns of activations, considering information distributed across all sensors simultaneously.

To compute a time-resolved Multivariate Pattern Analysis, a classification model is trained and cross-validated for each time point and participant individually, extracting different performance metrics according to the cfg structure (please see Figure 6). All of this process is coded in the function **mvpalab_mvpa(**cfg,fv), which computes the decoding analysis completely:

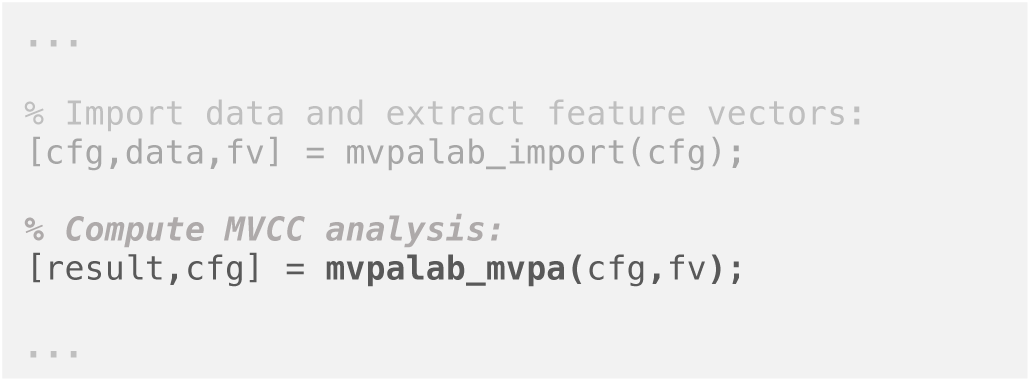

This function returns an updated version of the configuration structure (`cfg`) and the result variable (`result`). Performance values are stored in data matrices `[1 x time x subject]` inside the result variable as shown in the following figure:

For example, the time-resolved accuracy values can be extracted from result.**acc**. Other class-specific performance metrics such as f1-score, recall or precision are stored for each condition in:

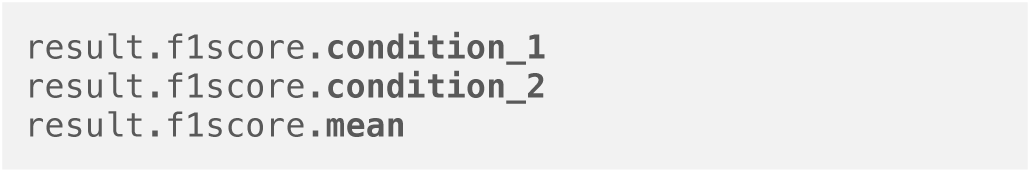

#### 2.4.2 TIME-RESOLVED MULTIVARIATE CROSS-CLASSIFICATION (TR-MVCC)

As mentioned before, the former MVPA technique has the ability to detect subtle differences in brain activation patterns. Thus, this powerful capacity could be used to study how these patterns are consistent across different cognitive contexts. In general, the consistency of the information across different sets of data can be analyzed with these techniques. To this end, classification models are trained with one set of data and the consistency is assessed by testing these models with another data sets, belonging to a different experimental condition. This technique is called Multivariate Cross-Classification (MVCC) [67] and is growing in popularity in recent years [68–70].

It is important to stress that different results can be obtained depending on which set is used for training and which one for testing (Train: **A** → Test: **B** or Train: **B** → Test: **A**). This is called classification direction. The observation of classification direction asymmetries in MVCC can be explained by several and very different phenomena, including complex neurocognitive mechanisms or a simple signal-to-noise ratio difference across datasets. For this reason, reporting results in both directions is highly recommended [71]. By default, MVPAlab computes and reports both directions separately. To compute the MVCC analysis, the function `mvpalab_mvcc` should be called after the feature extraction stage:

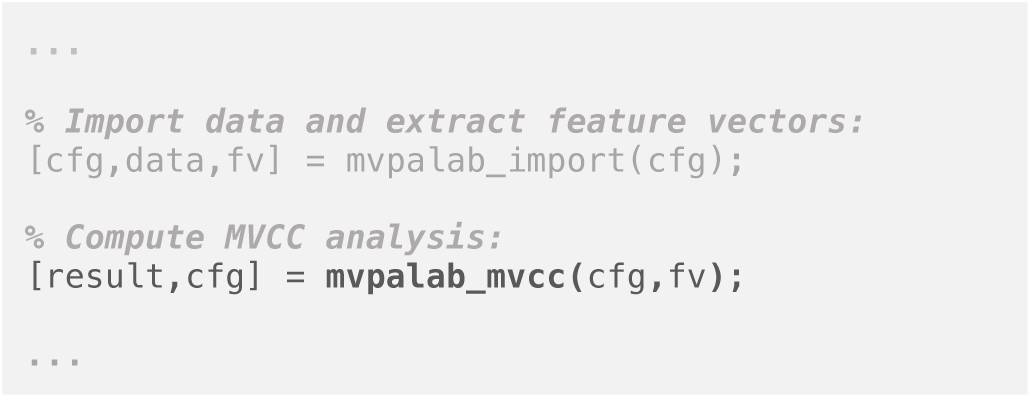

Similar to previous analysis, this function returns an updated version of the configuration structure and the results variable. In this case, time resolved accuracy values are stored for both classification directions in:

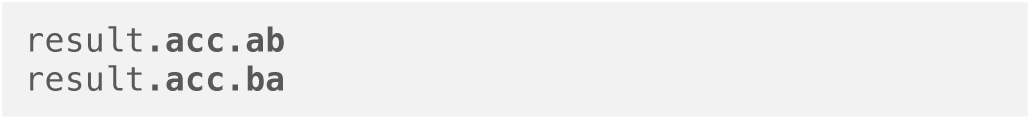

#### 2.4.3 TEMPORAL GENERALIZATION MATRIX

To evaluate the stability of brain patterns along time, temporal generalization analyses are commonly used. To obtain the temporal generalization matrix, the model is trained in a specific temporal point, testing its ability to discriminate between conditions in the whole temporal window. This process is then repeated for every timepoint thus obtaining the final decoding accuracy matrix (see Figure 8). An above-chance discrimination rate outside the diagonal of the matrix suggests that the same activity pattern is sustained in time. This phenomenon is usually interpreted as a reactivation of neural representations [66]. Therefore, if there is no evidence of temporal generalization, different patterns of activity can be inferred [57]. However, a recent study demonstrated that this interpretation is not always valid, suggesting that this phenomenon can be explained as an artefact of the manner in which the decoding accuracy provided by different components of the signal combine to bring about the overall decoding accuracy [72]. Regardless of the previously selected type of analysis (MVPA or MVCC), the calculation of the temporal generalization matrix can be enabled in the MVPAlab configuration structure as follows:

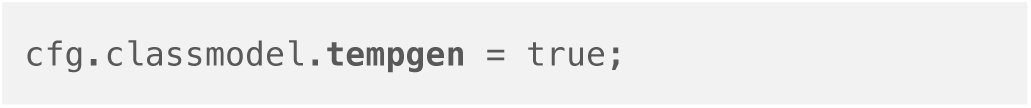

**Figure 8.**
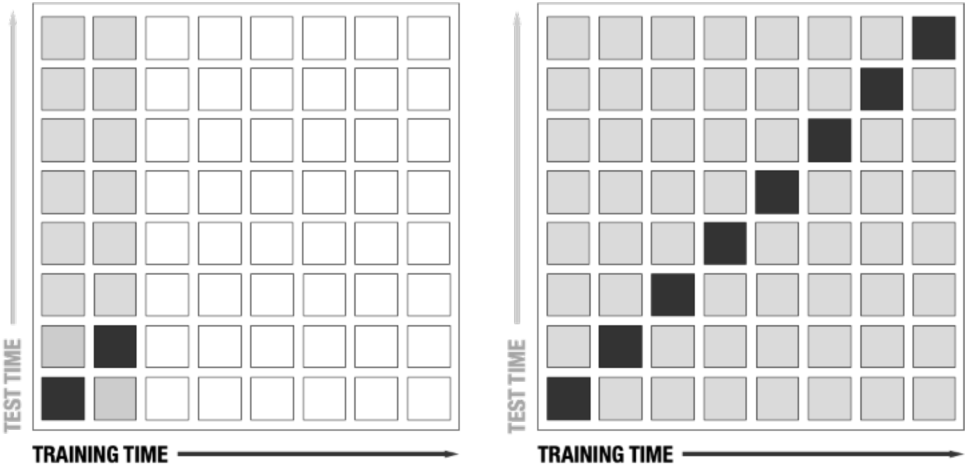
Temporal generalization matrix: the classification model is trained with data at certain time point (black square). This model is then tested along the remaining time points (grey square), repeating this process for each time point inside the epoch.

#### 2.4.4 FEATURE CONTRIBUTION ANALYSIS

Usually, classification algorithms are treated as black-boxes. However, highly useful information can be extracted out under specific circumstances. For example, the value of a feature weight, obtained after the training process of SVM models, is sometimes correctly interpreted as a measure of its contribution to the model decision boundary. In other words, it is a measure of its importance. As shown in Figure 3, the feature weight vector represents the coefficients of *ω*, which is an orthogonal vector to the separation hyperplane. However, as mentioned above, this is valid under certain scenarios (e.g. linear classifiers, use of the same scale for all features, no data transformations such PCA, etc.). Even meeting all these requirements, the interpretation of raw feature weights can lead to wrong conclusions regarding the origin of the neural signals of interest. A widespread misconception about features weights is that channels with large weights should be related to the experimental condition of interest, which is not always justified [73]. In fact, large weight amplitudes can be observed for channels not containing the signal of interest and *vice versa*. To solve this problem, Haufe et al. [73] proposed a procedure to transform these feature weights so they can be interpreted as origin of neural processes in space, which leads to more accurate predictions in neuroscience studies.

This useful procedure is implemented in the MVPAlab Toolbox. During any decoding analysis, MVPAlab extracts and saves the raw weight vectors and its Haufe correction in a time-resolved way. Thus, the contribution (or importance) of each electrode to the classification performance can be evaluated at any given timepoint. Additionally, and only if channel location information is available, MVPAlab can create animated plots representing the evolution of the distribution of weights over a scalp template. This analysis can be computed at group level or only for a specific participant. Please, see the *Result* section for further details.

Feature contribution analysis is disabled by default but can be enabled in the configuration file as follows:

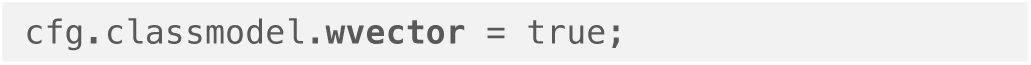

#### 2.4.5 FREQUENCY CONTRIBUTION ANALYSIS

The contribution of different frequency bands to the overall decoding performance can be assessed in MVPAlab through an exploratory sliding filter approach. To this end, the original EEG signal can be pre-filtered using a band stop sliding FIR filter. Therefore, different frequency bands can be filtered-out of the original EEG data, producing new filtered versions of the original dataset. The former time-resolved multivariate analysis is now computed for each filtered-out version of the data. The importance of each filtered-out band is quantified computing the difference maps in decoding performance between the filtered and the original decoding results. Accordingly, if the classification performance at any given point is higher for the original signal compared to the filtered-out version, then the removed frequency band contains relevant information used by the classification algorithms to discriminate between conditions. This procedure is illustrated in Figure 9. By definition, this analysis can be computed in a time-resolved manner (without temporal generalization) and using only the mean accuracy or the AUC as performance metric.

**Figure 9.**
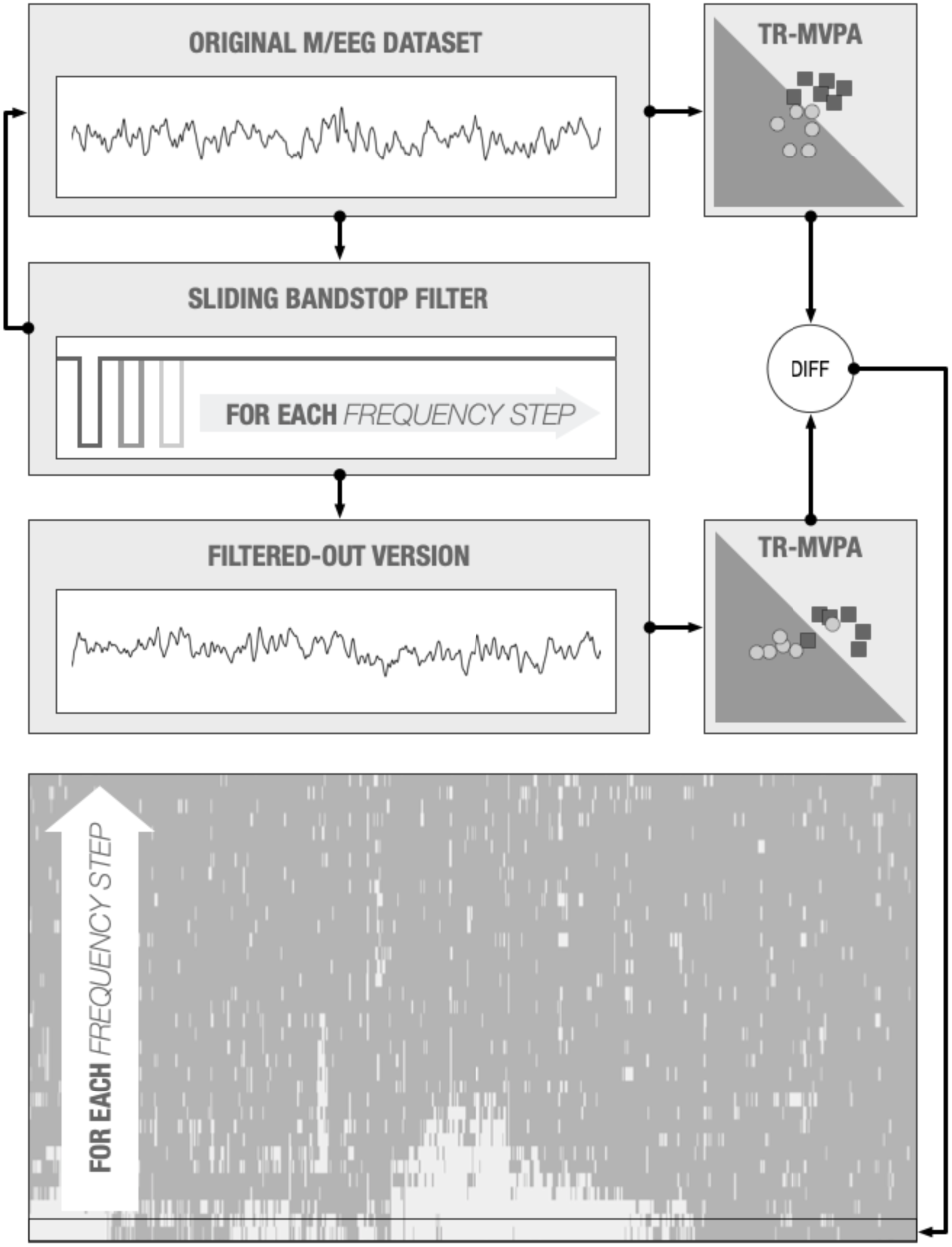
Sliding filter analysis diagram. This analysis compares in a time-resolved way the classification performance between the original dataset and a filtered-out version in which a certain frequency band has been removed. This procedure is repeated for each frequency band (step) returning a classification performance difference map which indicates how each frequency band contributes to the classification performance.

Several parameters should be defined in the MVPAlab configuration structure to compute the sliding filter procedure:

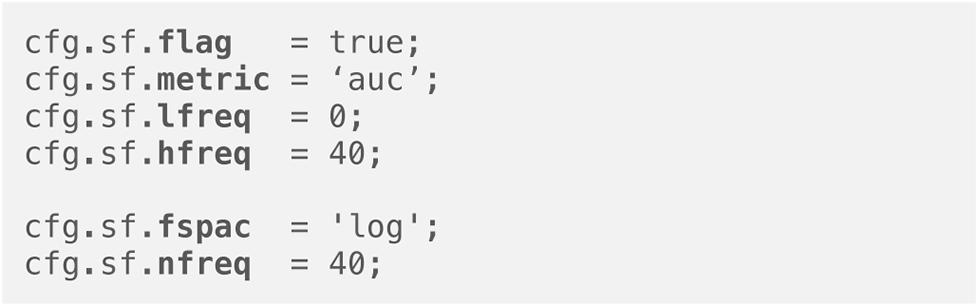

Sliding filter analysis can be enabled or disabled setting the configuration variable (.**flag**) to true or false. The (.**lfreq**) and (.**hfreq**) variables define the frequency limits in which the analysis will be computed. As mentioned before, mean accuracy (.**metric** = ‘acc) or AUC (.**metric** = ‘auc’) can be selected as performance metrics for this analysis. The number of individual frequency bands that will be removed from the original dataset (frequency resolution) is defined by (.**nfreq**). Each of these frequency bands can be linear (.**fspac** = ‘lin’) or logarithmically (.**fspac** = ‘log’) spaced as shown in Figure 10. On the one hand, if the frequency bands are linearly spaced, the frequency resolution is equally distributed across the entire spectrum. On the other hand, a higher frequency resolution is found in the low part of the spectrum if the frequency bands are logarithmically spaced. This is especially interesting for investigations focusing in the study of the lower part of the M/EEG spectrum (α, β and θ frequency bands).

**Figure 10.**
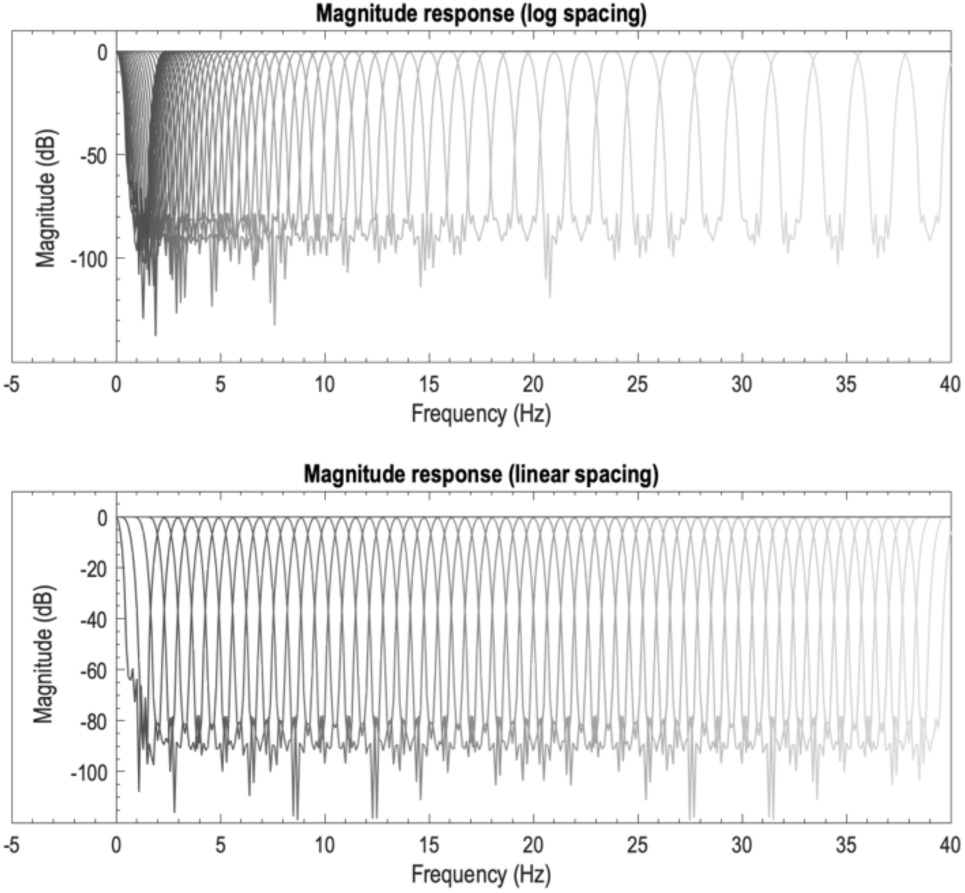
Removed frequencies: magnitude response for both linear and logarithmically spaced band-stop sliding filters. 60 frequency bands, 1408 filter order, Blackman window, 2Hz overlapped bandwidth.

The filter design parameters such as filter type (.`ftype`), filter bandwidth (.**bandwidth**), window type (.**wtype**), filter order (.**order**), and others, can also be tuned in the configuration file as follows:

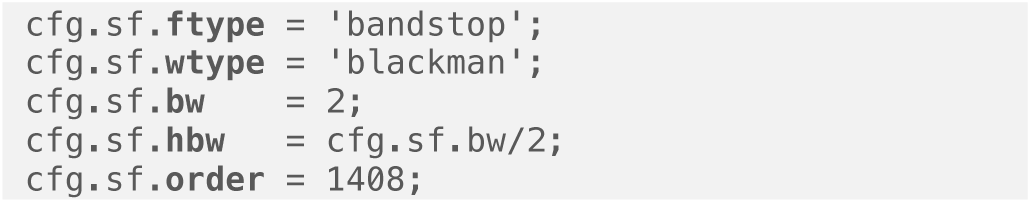

The filter design and filtering process employed by MVPAlab is based on the EEGlab built-in function *pop_firws*

Digital filters usually affect brain signals and are commonly applied at many stages from the data acquisition to the final publication. Many undesired events including temporal blurring or signal delays may occur, which may lead to incorrect interpretation of the results. Therefore, an appropriate filter design becomes crucial to prevent (or mitigate) these signal distortions. Please see [74,75] for a deeper understanding of how brain signals can be affected by filtering processes.

The complete sliding filter analysis pipeline is coded in both **mvpalab_import(**cfg) and **mvpalab_sfilter(**cfg) functions:

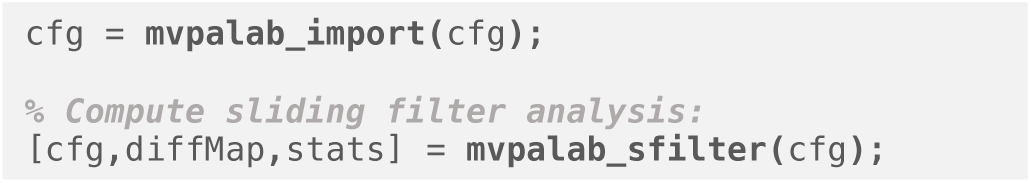

Due to the elevated RAM requirements of this analysis, the import function stores each filtered versions of the original dataset in a specific folder of your hard drive for each participant individually. The user should consider using an external hard drive for this high-demand analysis.

Then, as explained before, the function `mvpalab_sfilter()` computes and compares the decoding performance of different metrics between the original dataset and each filtered version, returning a difference map structure `diffMap`. The result matrices `[freqs x timepoints x subjects]` for specific performance metrics can be extracted using dot notation (e.g. diffMap.**auc**). Only the mean accuracy and the area under the curve are implemented for this analysis.

Additionally, if enabled, this function also implements the statistical permutation analysis, returning the `stats` variable, which includes the statistically significant clusters (Please see section 3.4 Statistical analysis for a detailed explanation).

### 2.5 Cluster-based permutation testing

In order to draw statistical inferences at the group level, MVPAlab implements a non-parametric cluster-based permutation approach, as proposed by Stelzer [76] for fMRI studies. This method has been adapted to electroencephalography data and can be computed for different performance metrics: mean accuracy, area under de curve, F1 score, recall and precision.

Using a combined permutation and bootstrapping technique, the null distribution of the empirical decoding accuracy is obtained. By default, at the single-subject level, 100 randomly permuted accuracy maps are generated. Then, one of the previously calculated accuracy maps for each participant is randomly drawn. This selection is group-averaged and the procedure is repeated 10^5^ times, generating 10^5^ permuted group accuracy maps. Next, for each timepoint, the chance distribution of accuracy values is estimated. The above and below chance thresholds are determined (99.9th percentile of the right and left-tailed area of the distribution), which correspond to a very low probability of obtaining significant results by chance (Figure 11). Then, clusters of time-points exceeding the previously calculated threshold in all the 10^5^ permuted accuracy maps are collected, generating the normalized null distribution of cluster sizes. Finally, a correction for multiple comparisons (False Discovery Rate (FDR)) is applied at a cluster level to obtain the smallest cluster size to be considered significant.

**Figure 11.**
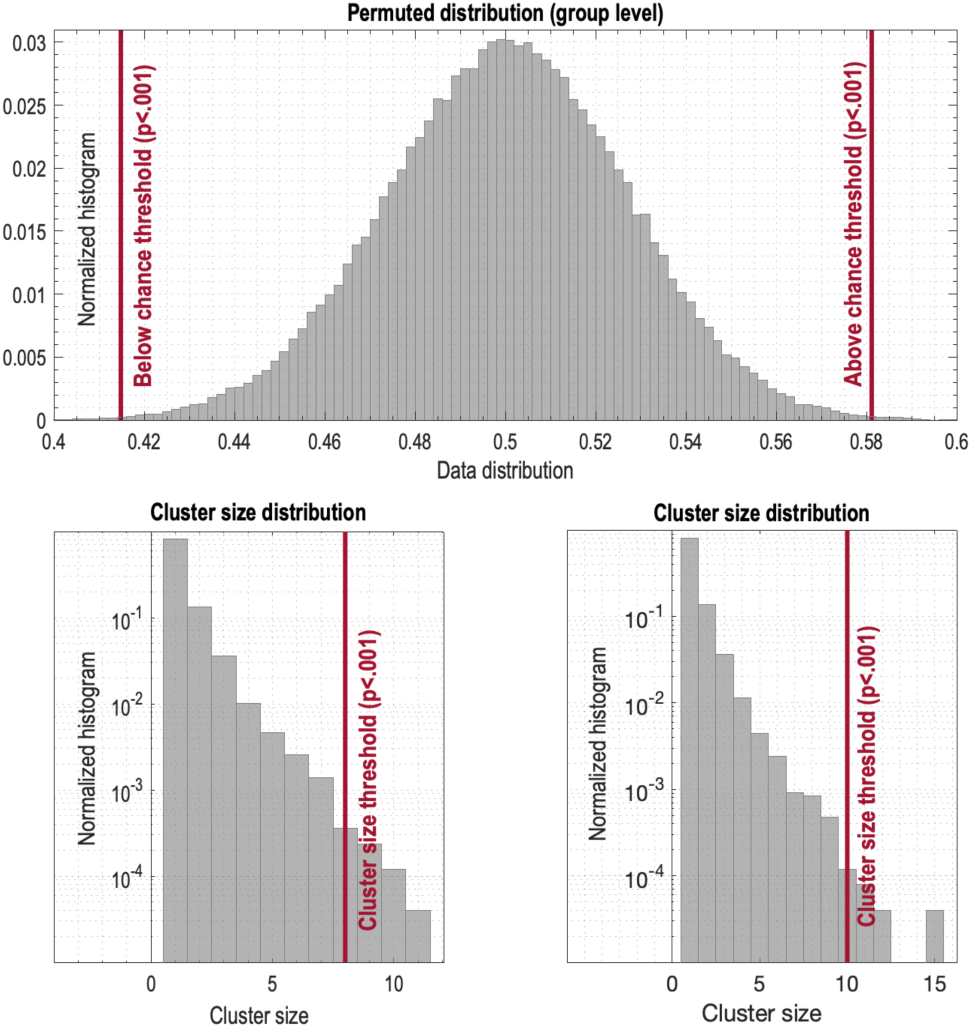
Accuracy and cluster size null distributions. The vertical line represents the threshold corresponding to a very low probability to obtain significant results by chance. These thresholds correspond to a p-value below 0.001 for both distributions.

The default parameters for this analysis can be modified in the MVPAlab configuration file as follows:

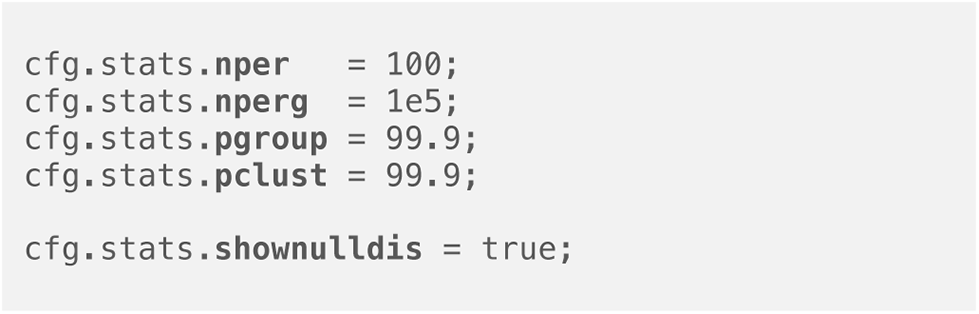

Two different functions coded the beforementioned pipeline:

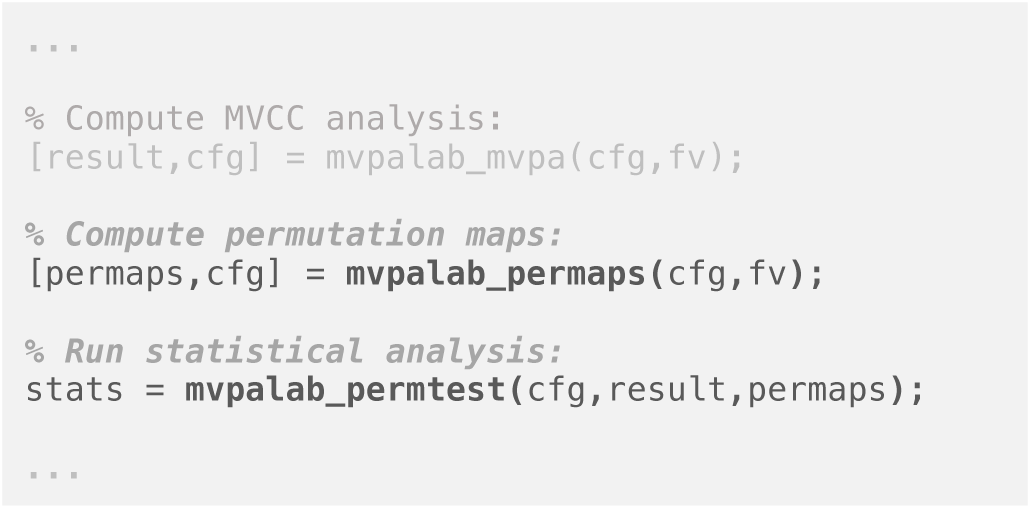

First, the function `mvpalab_permaps()` computes the required permuted accuracy maps for each subject, randomly shuffling the original class labels. Then, `mvpalab_permtest()` generates the null distributions, determines the significance thresholds, collects significant clusters, computes cluster size distributions and corrects for multiple comparisons (FDR) to obtain the smallest cluster size to be considered significant. The variable `stat` is returned containing, among others, below and above chance significant clusters:

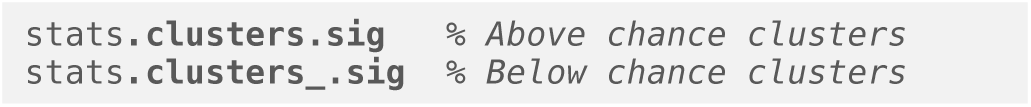

Above and below chance clusters are extracted using the MATLAB built-in function *bwconncomp* included in the Image Processing Toolbox.

### 2.6 Result representation pipeline

In addition to the graphic user interface, MVPAlab implements different high-level functions to generate highly-customizable graphical representation of the results. Once the decoding analysis is completed and the results files are saved, the graphical representation pipeline runs as follows:

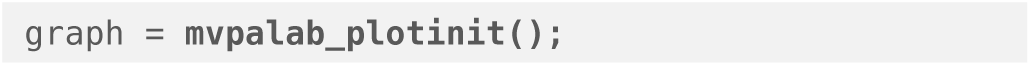

First, the function `mvpalab_plotinit()` generates and returns a default configuration structure (`graph`) containing all the required configuration parameters. Then, the specific result file to be plotted should be loaded:

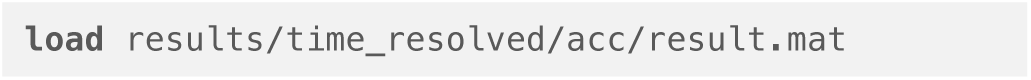

Finally, the high-level plotting function returns the graphical representation of the selected result file:

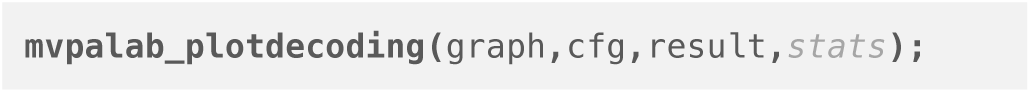

The variable `stats` is optional and contains, among others, the statistically significant clusters. If this variable is not omitted, significant results will be highlighted in the resulting figure. Several plotting functions are available for different types of analysis:

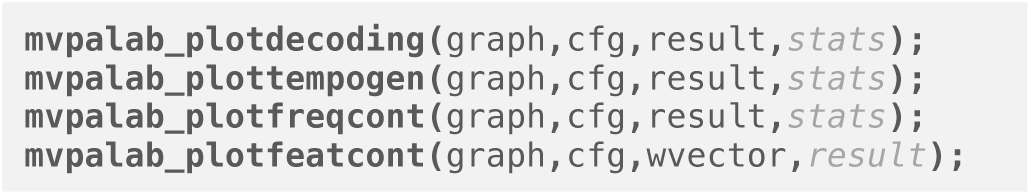

The `mvpalab_plotdecoding() function` generates time-resolved performance plots, `mvpalab_plottempogen()` is used for the graphical representation of temporal generalization matrices, `mvpalab_plotslidfilt()` function generates the graphical representation for the sliding filter analysis and `mvpalab_plotfeatcont()` can generate topological representations and temporal animations of features contribution to the decoding performance.

To get the best of the MVPAlab Toolbox plotting capabilities the use of the graphic user interface is highly recommended. This is a fast, flexible and very intuitive manner to design high-quality plots. Even so, the same results can be obtained by hand coding several configuration parameters included in the `graph` configuration structure. A complete selection of the most useful configuration parameters and a short explanation is listed below:

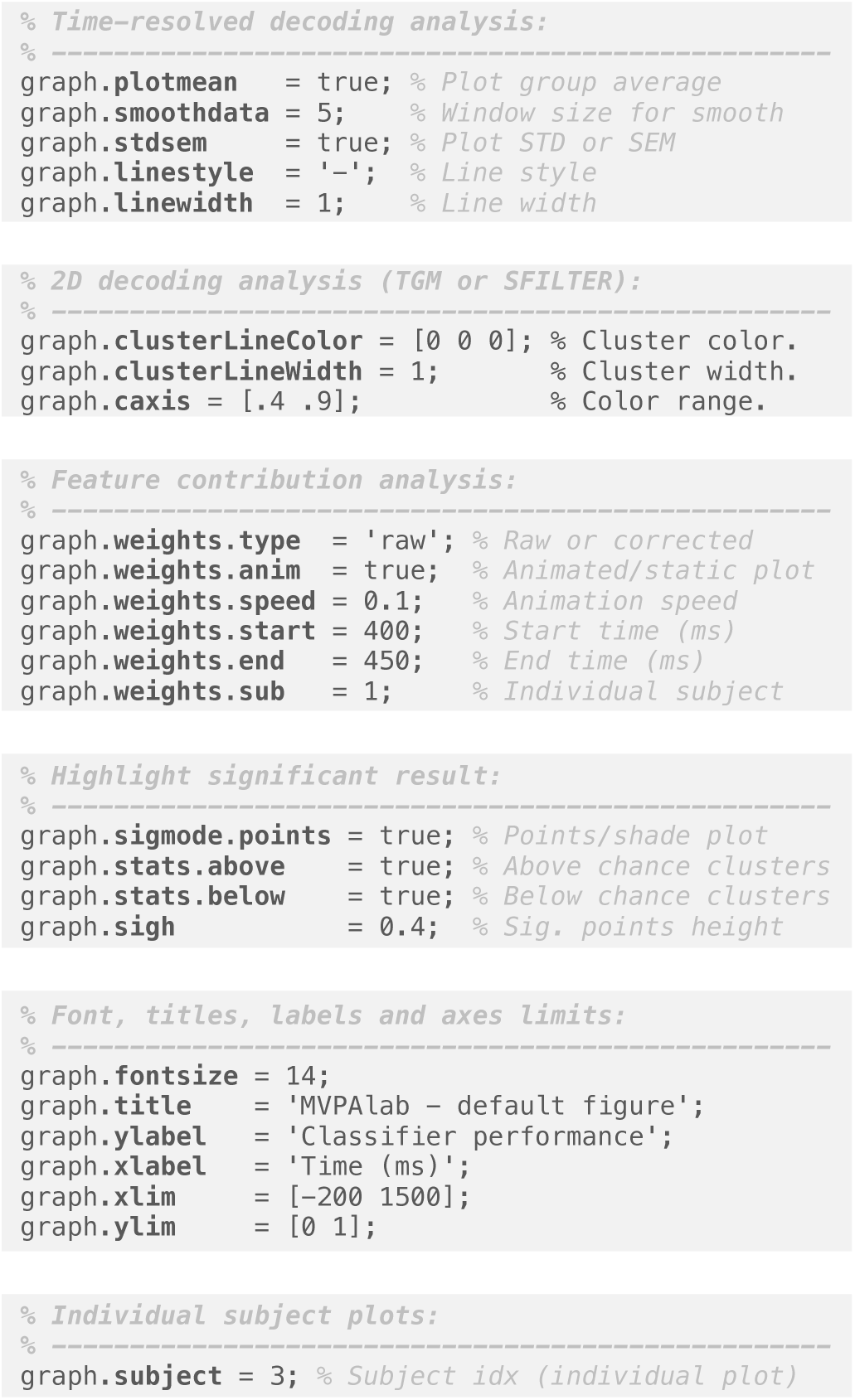

Finally, for a correct visualization of the results, ten new color gradients and colormaps (Figure 12) have been designed and incorporated to the MATLAB predefined ones. The default MATLAB colormap can be modified as follows:

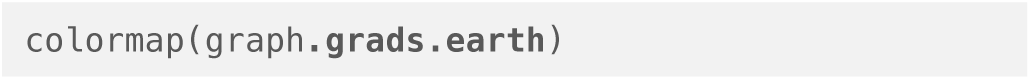

**Figure 12.**
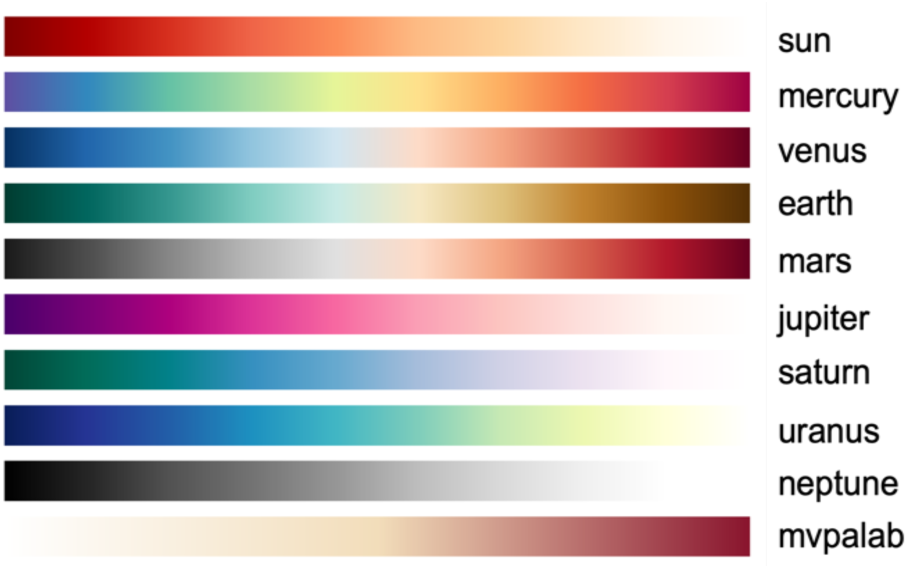
Ten color maps included in MVPAlab Toolbox.

## 3. RESULTS

During this section we present the results obtained after testing all the MVPAlab main functionalities with the sample EEG dataset presented in Section 2 *Materials and Methods*. As mentioned, we compiled this sample dataset for illustration purposes, including the EEG data of two main conditions (or classes) and four subconditions of three different participants. Readers interested on the results obtained for the entire sample should refer to the original publication [44].

### Time-resolved decoding analysis

**Figure 13 (a)** depicts the result of a time-resolved decoding analysis comparing the classification performance of two models, linear support vector machine and linear discriminant analysis. Shaded areas represent the Standard Error of the Mean (SEM) of the averaged performance across participants. Additionally, single-subject plots are depicted in dashed and dotted lines. Statistically significant areas for each classification model are highlighted using horizontal color bars. As shown, SVM outperforms LDA by obtaining higher performance and a wider significant window.

**Figure 13.**
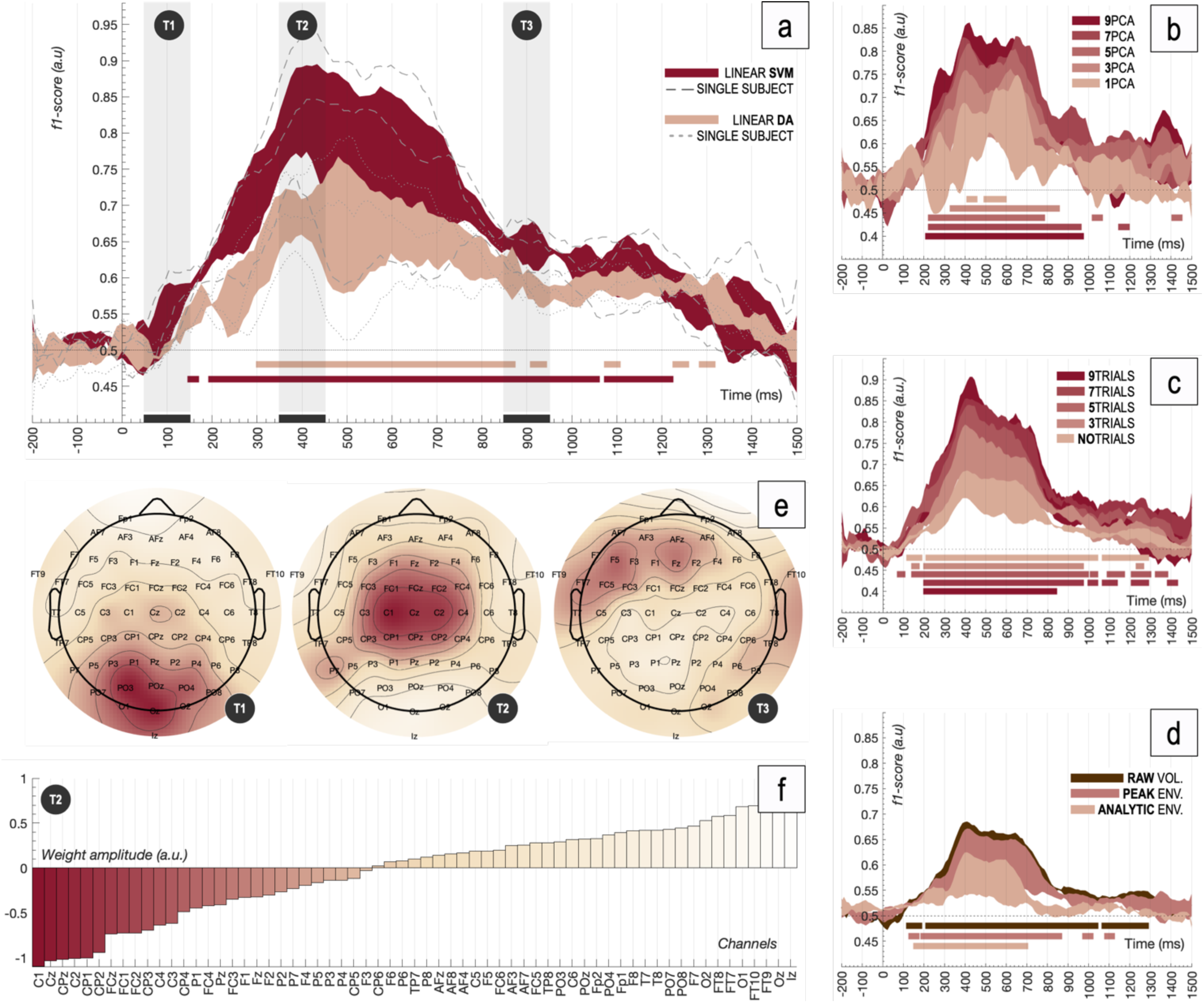
Time-resolved MVPA results. **(a)** Decoding performance (f1-score) for different classification models at a group-level: support vector machine vs. linear discriminant analysis. Single subject plots are represented in dashed and dotted lines. Significant clusters are highlighted using horizontal colored bars. Shaded areas represent the standard error of the mean. **(b)** Group-level decoding performance for different number of features when PCA is applied. **(c)** Group-level decoding performance as a function of the selected number of trials to average. **(d)** Group-level decoding performance when different power envelopes are extracted and employed as features instead of the raw voltage. **(e)** Group-level weight distribution (corrected) for three different time windows: T1: 50-150ms, T2: 350-450ms and T3: 850-950ms. **(f)** Weights’ amplitude for each channel sorted by importance.

To compute this MVPA analysis, classification models were trained using smoothed (5 timepoint moving average) and normalized supertrials (8 trials randomly averaged). No PCA was computed, so raw voltage values were extracted from the 64 electrodes as features in balanced datasets.

### Dimensionality reduction

**Figure 13 (b)** shows the time-resolved classification performance (f1-score) averaged across participants of an SVM classifier, using different number of PCA components as features. As shown, the f1-score increases with the number of features. Significant results were obtained employing just the first PCA component. When only the first nine PCA components were employed as features, the classification model showed comparable performance results to those obtained when no PCA is computed, as depicted in Figure 13 (a). Computation time is in fact reduced when the dimension of the feature space is smaller, however, when PCA transformation is computed, the original spatial information is lost.

### Supertrial generation

**Figure 13 (c)** depicts the classification performance when the input dataset was reduced by randomly averaging different numbers of trials belonging to the same condition. This trial averaging process generates supertrials with an increased signal-to-noise ratio. As shown, the SVM model performance increases with the number of trials averaged, however, the variability of the data (the standard error of the mean) also does due to the reduced input dataset. Thus, this figure shows wider significant windows when no or few trials are averaged.

### Power envelope as feature

The comparison between the performance of classifiers using different EEG signal characteristics as features is showed in **Figure 13 (d)**. First, the peak and analytic upper envelopes of the EEG signal were calculated (5 timepoints window). Then, feature vectors were extracted from these power signals. Significantly lower performance rates were obtained for the analytic power envelope. Although the main goal of this article is not to address this type of questions, there seems to be a plausible cause favoring this outcome: the phase of the EEG signa may contain critical information to discriminate between the two experimental conditions employed. This is due to the fact that the instantaneous phase information contained within the original EEG signal is discarded during the analytic power envelope computation (see *Appendix B* for further details). This approximation is employed in recent literature [72][77] to remove instantaneous phase from certain brain oscillations and to study how this phase information contributes to decoding performance.

### Feature contribution analysis

During the training process of the previous linear SVM model, the feature weights were calculated for each timepoint and subject and corrected according to Haufe’s method [73]. In order to show the activity distribution contributing to decoding accuracy, the feature weights were averaged across participants and three different temporal windows. First, when the slope of the decoding curve becomes positive, between 50-150ms. Then, between 350-450ms, when decoding performance peaks, and finally between 850-950ms, at the end of the significant window for LDA. A corrected version of the training weights distribution for these three different time windows is depicted in **Figure 13 (e)**. Finally, **Figure 13 (f)** shows the weight amplitude of each channel sorted by its importance, averaged across participants during the 350-450ms temporal window.

### Temporal generalization analysis

**Figure 14 (a)** shows the temporal generalization matrix of the first MVPA analysis, Figure 13 (a), representing the performance value (AUC) for each combination of training-test time points. Above-chance significant clusters are highlighted using black lines. This approach is an extension of time-resolved decoding, which is an indication of how EEG patterns vary or persist in time. Different performance metrics, such as the area under the curve or the mean accuracy are usually reported, generating temporal generalization patterns that resembles those shown in Figure 14 (a). This way, above-chance performance clusters outside the diagonal of the matrix are interpreted as a sign of temporal stability of certain activity patterns along time.

**Figure 14.**
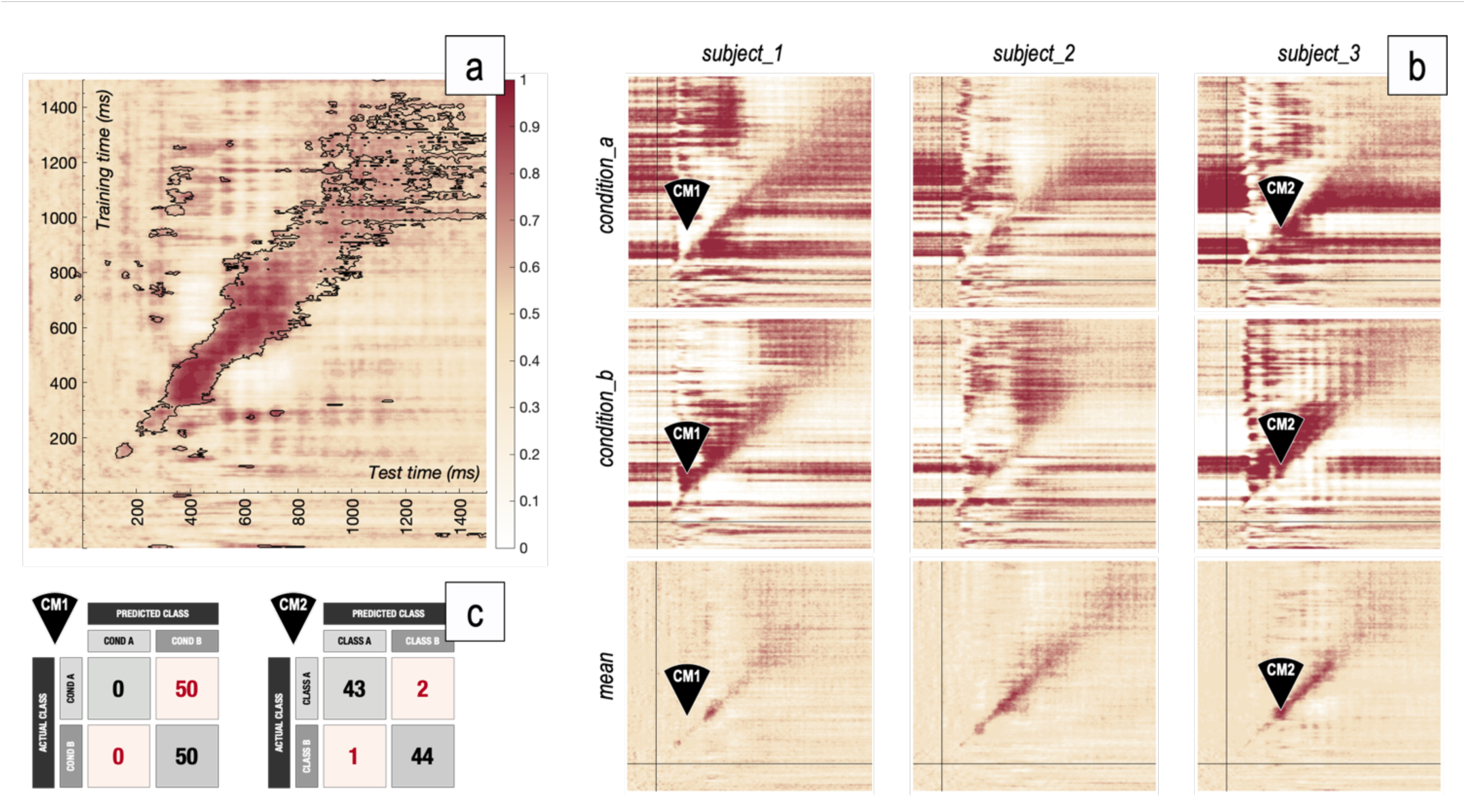
Temporal generalization results. **(a)** Group-level temporal generalization matrix (area under the ROC curve) for an SVM classifier when 8 trials were averaged to generate the input dataset. Above-chance significant clusters are highlighted using black lines. **(b)** Single subject generalization patterns (sensitivity), individually calculated for each condition. **(c)** Confusion matrices CM1 and CM2 for two different timepoints marked in (b).

However, in-depth examinations revealed interesting behaviors of classification models, providing extra information about how individual conditions are classified, especially in those areas in which no temporal generalization occurs. **Figure 14 (b)** depicts the sensitivity (recall) of the classification model for each condition and subject. Complementary generalization patterns are observed for individual conditions, revealing extreme sensitivity values especially when no temporal generalization occurs. Some examples are presented and analyzed using the corresponding confusion matrices. As seen in **Figure 14 (c)**, the confusion matrix **CM1** indicates that, for this specific temporal point, no test samples belonging to *condition_a* were correctly predicted as *condition_a*, leading to a sensitivity value of 0 for this condition. By contrast, all samples belonging to *condition_b* were correctly labelled (in addition to all samples belonging to *condition_a* incorrectly predicted as *condition_b)* which leads to a sensitivity value of 1. This behavior is frequent across subjects and timepoints, reflecting the inability of the classifier to correctly predict information in several areas, which is a clear sign of the absence of temporal persistence of patterns.

### Multivariate Cross-Classification analysis

**Figure 15 (a)** depicts the result of a time-resolved multivariate cross-classification analysis. The classification model was trained with *condition_1* vs. *condition_2* and *condition_3* vs. *condition_4* were used for testing. This process was repeated inversely, generating two different decoding performance curves corresponding to both classification directions (train: A, test: B and *vice versa*). Additionally, single subject curves were added to the figure for each classification direction. As shown, windows of significant differences are obtained between 200-800ms, indicating that this technique successfully shows the consistency of patterns across different sets of data.

**Figure 15.**
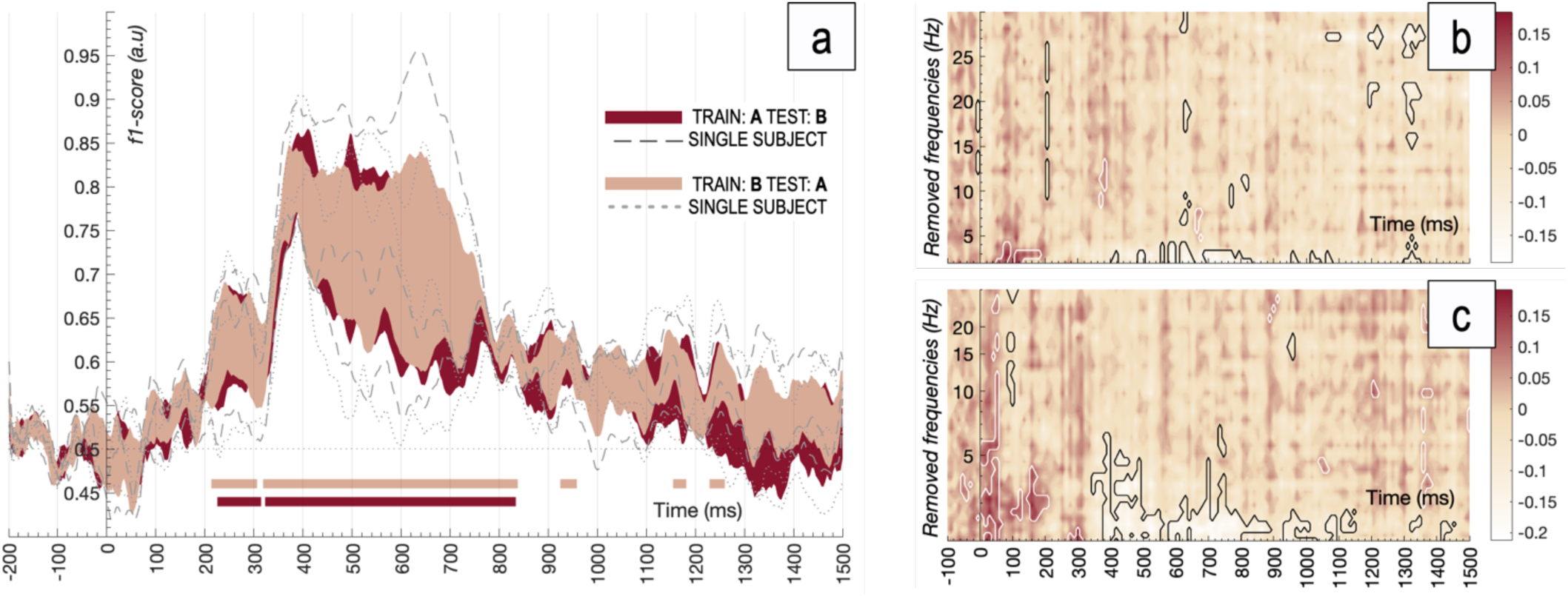
Time-resolved MVCC and frequency contribution analysis results. **(a)** Group-level decoding performance (F1-score) for both cross-classification directions. Single subject plots are represented in dashed and dotted lines. Significant clusters are highlighted using horizontal colored bars. Shaded areas represent the standard error of the mean. **(b-c)** Decoding performance maps when different frequency bands are removed from the original datasets in a linear and logarithmically spaced steps.

### Frequency contribution analysis

A sliding band-stop filter approach was followed to study the contribution of each frequency band to the overall decoding accuracy. The band-stop FIR filter was designed using the EEGLAB `pop_firws` function (2Hz bandwidth, 0.2Hz transition band, 2048 filter order, Blackman window). The original EEG dataset was pre-filtered (32 overlapped frequency bands, between 0–30Hz in linear and logarithmically-spaced steps) producing 32 new filtered versions of the original signals. The former time-resolved decoding analysis (*condition_a* vs. *condition_b*) was conducted for each filtered version and the importance of each filtered-out band was quantified computing the difference maps in decoding performance between the filtered and the original decoding results. **Figures 15 (b)** and **(c)** show the results of the sliding filter analysis for linear and logarithmically-spaced steps respectively. As shown, decoding accuracy significantly dropped when frequencies up to 6Hz were filtered-out, suggesting that the studied phenomenon relies on processes operating in the Delta and Theta frequency bands. Significant clusters were calculated applying the proposed cluster-based permutation test to filtered-out datasets, generating accuracy null distributions for each time-frequency point.

## 4. DISCUSSION

Despite the MVPAlab Toolbox is freely available, an important limitation is that it needs the MATLAB core to be executed, which is a proprietary and expensive software. We are aware of the recent growth of free software alternatives, such Python, in academic environments. Nevertheless, we built this software under MATLAB due several reasons, including the huge amount of available and well-documented functionalities for this platform, their active user community and its wide implementation in neuroscience labs. Even so, there are excellent open source alternatives for those users with no access to a MATLAB license.

Additionally, the MVPAlab Toolbox is not yet compatible with BIDS-EEG [78] format, which is a recently developed project for electroencephalography studies, extending the original Brain Imaging Data Structure [79] (BIDS). Both projects are an excellent effort to standardize the way data is stored, increasing accessibility, usability and reproducibility of neuroimaging data. We favor these principles and we are planning to integrate BIDS-EEG format in the MVPAlab Toolbox in future releases.

Classification algorithms are the cornerstone of multivariate decoding analyses. However, these powerful techniques suffer from hyperparameter overfitting, which usually leads to invalid result. A recent study refers to this phenomenon as “*overhyping*”[80] and proposes several strategies to avoid this problem. Regular cross-validation approaches are commonly employed to mitigate spurious result in classification accuracies, but it has been proved that, in some cases, they are not sufficient [80]. Several strategies, such as pre-registration, nested cross-validation [81], lock box and blind analyses are presented as reliable alternatives to prevent or mitigate *overhyping*. Unfortunately, the MVPAlab toolbox does not currently implement those strategies, but we are further investigating these issues for future releases. Additionally, recent studies [82,83] proposes the Statistical Agnostic Mapping (SAM) as an interesting alternative to the cross-validation procedures. Particularly in neuroscience, these approaches usually leads to small sample sizes and high levels of heterogeneity when conditions are split into each fold, causing among other things, a large classification variability [84]. To address these problems, SAM considered the use of the resubstitution error estimate as a measure of decoding performance. The difference between the actual error and the resubstitution error (which is a very optimistic measure) is upper-bounded by a novel analytic expression proposed in the original article. See [82,83] for further details. Future releases of the MVPAlab Toolbox are planned to include this novel classification paradigm, which at the moment is under development.

Furthermore, dimensionality reduction is a crucial step in neuroimaging studies to select the most relevant predictor variables, reducing the experimental noise and mitigating the *small-n-large-p* problem. These techniques prevent the classification model from overfitting, leading to a better predictions and increasing its generalization capability [85]. Although MVPAlab implements Principal Component Analysis, which is one of the most popular dimensionality reduction approaches in neuroscience studies, there are different algorithms which have not been implemented yet. The integration with some of these feature reduction approaches, such as Partial Least Square (PLS)[86], is currently under development. Regarding to the classification stage, the MVPAlab Toolbox implements probably two of the most commonly employed classification algorithms in neuroscience literature: Support Vector Machines and Discriminant Analysis, in their linear and non-linear versions. However, this configuration may not be enough in certain situations. In fact, different software alternatives include many other classification models, such as Logistic Regressions, Naïve Bayes or ensembles methods. As mentioned, the MVPAlab Toolbox is in constant development, these functionalities are planned to be implemented in near future.

The MVPAlab Toolbox was initially developed for M/EEG analysis. Due to its nature, M/EEG signals provide exceptional temporal resolution, but lack spatial resolution. Contrary, other non-invasive techniques, such as fMRI, can identify brain activity changes at millimetric levels but suffer from poor temporal resolution. To overcome this dichotomy, recent trends in the neuroimaging field opt for the multimodal data fusion[87,88], which is a step forward towards a better understanding of brain function. These fusion approaches combine data from different neuroimaging techniques (M/EEG-fMRI), preserving their strengths while overcoming their weaknesses[89]. Extending the MVPAlab functionality from multivariate M/EEG analyses to multimodal data fusion represents one of the most important lines of development on the MVPAlab roadmap.

There are a myriad of new analyses and techniques that can be employed to analyze data of different nature in neuroscience, which is a clear indicator of the fast growth of the field. As mentioned, MVPAlab Toolbox was initially designed to work with epoched M/EEG data, extracting the raw potential of the signal and computing time-resolved classification analyses. The latest release supports different signal characteristics as features, such as the power envelope or the instantaneous phase of the signal. Recent studies [90] implement different feature engineering techniques, concatenating data from different frequency bands, to improve the classification result. Currently, MVPAlab does not implement these strategies. However, MVPAlab can be used as a general-purpose classification tool. Users can adapt and import their own datasets, regardless of its nature (source space data, connectivity data or not even M/EEG related signals), and easily perform time-resolved classification analyses.

## 5. CONCLUSIONS

MVPAlab is a very flexible, powerful and easy-to-use decoding toolbox for multi-dimensional electroencephalography data, including an intuitive Graphic User Interface for creation, configuration, and execution of different decoding analysis. Not a single line of code is needed. For those users with more coding experience, MVPAlab implements high and low-level routines to design custom projects in a highly flexible manner. Different preprocessing routines, classification models and several decoding and cross-decoding analyses can be easily configured and executed. MVPAlab also implements exclusive analyses and functionalities, such as parallel computation, significantly reducing the execution time, or frequency contribution analyses, which studies how relevant information is coded across different frequency bands. MVPAlab also includes a flexible data representation utility, which generates ready-to-publish data representations and temporal animations. All of this combined makes MVPAlab Toolbox a compelling option for a wide range of users.

## CODE VERSION AND AVAILABILITY

An up-to-date version of the toolbox is freely available in the following GitHub repository:

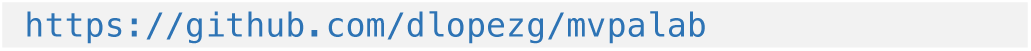

We use semantic versioning (e.g. X.Y.Z) to denote different releases, the most recent being the v1.0.0 version, which is our first public release including a stable version of the toolbox.

The software documentation can also be found in our GitHub repository:

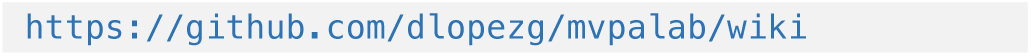

MVPAlab toolbox is released under a GNU General Public License (GPL) v.3.0, which allows users to freely use, change and share this software. For further license details please see:

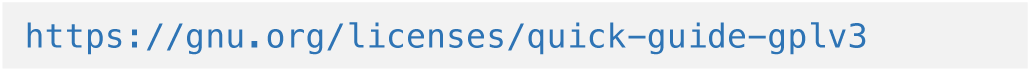

We encourage all users to collaborate in MVPAlab Toolbox development by submitting their own contributions and improvements via *pull request*. To suggest new features, bug report or any other related issues, please use the MVPAlab issue tracker available in GitHub in the following link:

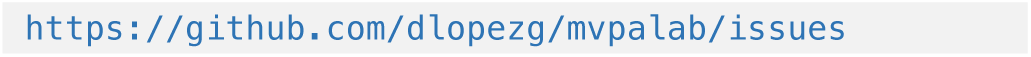

The sample EEG dataset used in this article is hosted in the Open Science Framework project:

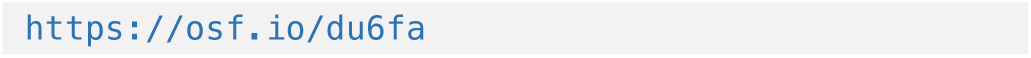

## SUPPLEMENTARY MATERIAL

Supplementary material such as temporal animations of feature contributions are available to download from an Open Science Framework project:

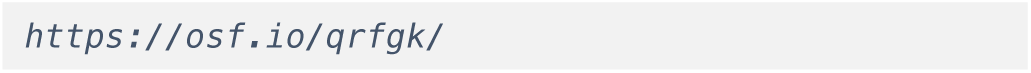

The Supplementary Material folder includes different video files (.mov) recording the temporal distribution of channels contributing to the decoding accuracy. Raw and corrected feature weights animations for individual participants and group-averaged are included.

## ACKNOWLEDGEMENTS

This research was supported by the Spanish Ministry of Science and Innovation under the TEC2015-64718-R and PID2019-111187GB-I00 grants. The first author of this work is supported by a scholarship from the Spanish Ministry of Science and Innovation (BES-2017-079769). The sample EEG dataset was extracted from an original experiment previously approved by the Ethics Committee of the University of Granada.

## Appendix A. Benchmarks and processing time

The performance comparison between different implementations of several classification libraries is out of the scope of this article. However, processing time for different analysis have been measured in Windows and macOS and are reported in the following table:

**Table 2.** Processing time in seconds for different task and platforms.

| Time (s) | Windows 10 (64 bits) |  | MacOS 11.3 (64 bits) |  |
| --- | --- | --- | --- | --- |
|  | Single | Parallel | Single | Parallel |
| **T1: TR-SVM** | 15.58 | **4.03** | 15.18 | **5.27** |
| **T1: TR-LDA** | 8.63 | **1.95** | 10.24 | **3.04** |
| **T1: TG-SVM** | 120.88 | **21.70** | 102.70 | **26.42** |
| **T1: TG-LDA** | 302.72 | **58.79** | 279.34 | **92.37** |
| **T2: TR-SVM** | 10.73 | **2.28** | 10.30 | **4.04** |
| **T2: TR-LDA** | 3.80 | **1.03** | 4.08 | **1.43** |
| **T2: TG-SVM** | 53.24 | **11.48** | 49.98 | **16.43** |
| **T2: TG-LDA** | 155.69 | **25.77** | 127.61 | **38.49** |

*Task 1 (T1)* consist of a single subject time-resolved decoding analysis and a five-fold cross validation stage, when only the mean accuracy was calculated, ten trial averaging and no dimensionality reduction was computed. In this scenario, different classification algorithms (SVM and LDA) were trained and validated for 256×256 timepoints using 80 observations (trials) and 63 features (electrodes).

*Task 2 (T2)* consist of a single subject time-resolved cross-decoding analysis, when only the mean accuracy was calculated, five trial averaging and no dimensionality reduction was computed. Both classification directions were calculated. In this scenario, different classification algorithms (SVM and LDA) were trained and validated for 256×256 timepoints using 80 observations (trials) and 63 features (electrodes).

These tests were computed in two different setups. First, in a 6-Core workstation (Intel Core i7-5820K CPU @ 3.30GHz, 32GB RAM DDR4 @ 2400MHz) running Windows 10 (64 bits) and MATLAB 2020a (9.8.0.1323502) and finally in a cuad-core MacBook Pro (Intel Core i7-6820HQ CPU at 2,7GHz, 16GB RAM LPDDR3 @ 2133MHz) running macOS Big Sur (64 bits, version 11.3) and MATLAB 2020a (9.8.0.1323502).

## Appendix B. Power envelope and instantaneous phase calculation

Different signal characteristics, such the instantaneous amplitude or phase, can be easily calculated and extracted in the complex plane. In order to extract this information from a real-valued signal *x(t)* (e.g. the electroencephalogram), the following transformation can be applied:

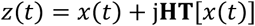

Here, *z(t)* is the complex form of *x(t)*, also known as the ‘*analytic signal*’, and **HT** denotes the Hilbert’s Transformation of the real-valued signal, defined as:

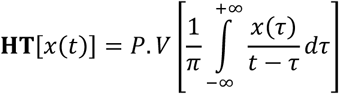

where P.V denote the *Cauchy Principal Value* of the integral, which is required for assigning values to improper integrals values that would otherwise be undefined (the singularity occurs when t = *τ*). Thus, the instantaneous amplitude, also known as power envelope *e*(*t*), or the instantaneous phase *ϕ*(*t*), can be easily extracted from the analytic signal as follows:

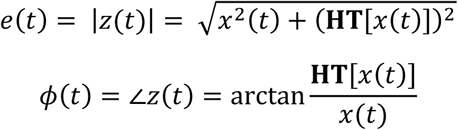

